# Towards Reproducible Brain-Wide Association Studies

**DOI:** 10.1101/2020.08.21.257758

**Authors:** Scott Marek, Brenden Tervo-Clemmens, Finnegan J. Calabro, David F. Montez, Benjamin P. Kay, Alexander S. Hatoum, Meghan Rose Donohue, William Foran, Ryland L. Miller, Eric Feczko, Oscar Miranda-Dominguez, Alice M. Graham, Eric A. Earl, Anders J. Perrone, Michaela Cordova, Olivia Doyle, Lucille A. Moore, Greg Conan, Johnny Uriarte, Kathy Snider, Angela Tam, Jianzhong Chen, Dillan J. Newbold, Annie Zheng, Nicole A. Seider, Andrew N. Van, Timothy O. Laumann, Wesley K. Thompson, Deanna J. Greene, Steven E. Petersen, Thomas E. Nichols, B.T. Thomas Yeo, Deanna M. Barch, Hugh Garavan, Beatriz Luna, Damien A. Fair, Nico U.F. Dosenbach

## Abstract

Magnetic resonance imaging (MRI) continues to drive many important neuroscientific advances. However, progress in uncovering reproducible associations between individual differences in brain structure/function and behavioral phenotypes (e.g., cognition, mental health) may have been undermined by typical neuroimaging sample sizes (median N=25)^1,2^. Leveraging the Adolescent Brain Cognitive Development (ABCD) Study^3^ (N=11,878), we estimated the effect sizes and reproducibility of these brain-wide associations studies (BWAS) as a function of sample size. The very largest, replicable brain-wide associations for univariate and multivariate methods were r=0.14 and r=0.34, respectively. In smaller samples, typical for brain-wide association studies (BWAS), irreproducible, inflated effect sizes were ubiquitous, no matter the method (univariate, multivariate). Until sample sizes started to approach consortium-levels, BWAS were underpowered and statistical errors assured. Multiple factors contribute to replication failures^4–6^; here, we show that the pairing of small brain-behavioral phenotype effect sizes with sampling variability is a key element in wide-spread BWAS replication failure. Brain-behavioral phenotype associations stabilize and become more reproducible with sample sizes of N⪆2,000. While investigator-initiated brain-behavior research continues to generate hypotheses and propel innovation, large consortia are needed to usher in a new era of reproducible human brain-wide association studies.

## Main

The advent of MRI has given us the remarkable ability to non-invasively map human brain structure^7,8^ (e.g. cortical thickness) and function (e.g. resting-state functional connectivity [RSFC])^9,10^. In addition to brain mapping, linking individual differences in brain structure and function to typical variation in behavioral phenotypes (e.g., cognitive ability, psychopathology) is a central goal of human neuroscience. Such brain-wide association studies (BWAS) hold great promise for predicting and reducing psychiatric disease burden and advancing our understanding of the cognitive abilities that underlie humanity’s intellectual feats. However, obtaining MRI data remains very expensive (~$1,000/hr), resulting in many small-sample BWAS studies (e.g., median N=25^1,2^), whose results often fail to replicate^1,11–15^.

Factors contributing to replication failures include methodological variability^6^, data mining for “significant” results (p-hacking), confirmation and publication biases^16^, and inadequate statistical power^4,5^. These factors of poor reproducibility are known from population-based research in psychology^17^, genomics (i.e. genome-wide association studies [GWAS])^18^, and medicine^19^. Behavioral and neuroimaging researchers are starting to address replication failures by standardizing analyses, pre-registering hypotheses, publishing null results, and data/code sharing^1^. While these open science efforts are improving reproducibility, reliance on sample sizes achievable by individual research groups may be the most significant obstacle to identifying reproducible brain-wide associations. If brain-wide associations have subtle true effect sizes (e.g., r=0.1), larger samples will be required to reproducibly measure them, simply due sampling variability (i.e., random variation). Sampling variability decreases and effect sizes stabilize towards their true values^20^ with increasing sample sizes, at the rate of 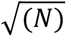. Therefore, epidemiology and genomics researchers were forced to steadily increase sample sizes from N<100 to N>1,000,000^18,21,22^ to overcome sampling variability and small effect sizes.

Recently, neuroimaging consortia aimed at linking brain measures to behavioral phenotypes have collected samples orders of magnitude larger than before, harmonizing imaging methodologies across sites (e.g., ABCD Study^3^, N=11,878; Human Connectome Project^23^ [HCP], N=1,200). Growing sample sizes have also facilitated the use of multivariate techniques (e.g., support vector regression [SVR], canonical correlation analysis [CCA]), with the hope of detecting reproducible brain patterns associated with behavioral phenotypes. Here, using the ABCD Study data, we performed a series of univariate and multivariate analyses to generate more precise estimates of brain-behavioral phenotype associations and to evaluate reproducibility as a function of sample size, ranging from typical (N=25) to very large (N=3,928).

### Brain-Wide Associations Cannot Be Estimated Precisely in Typically Sized Studies

Associations between the brain and behavioral phenotypes are classically estimated using univariate models in which a brain feature (e.g., RSFC between two regions [“edge”]) is correlated with typical variation in a behavioral phenotype (e.g., cognitive ability). To estimate the effect sizes of brain-wide associations in ABCD data, we correlated widely used cortical thickness and RSFC metrics with 41 measures indexing demographics, cognition, and mental health (Extended Data Table S1). Brain-wide associations were estimated across multiple levels of anatomical resolution in both structural (i.e., cortical vertices, regions of interest [“ROI”], networks; see Methods) and functional (resting-state connections [“edges”], principal components [“components”], networks) data. To ameliorate the effects of nuisance variables such as head motion, we applied extremely rigorous denoising strategies (N=3,928; >8mins RSFC data post frame censoring at a framewise displacement [FD] filtered<0.08mm). Repeat analyses employing less rigorous motion censoring, and thus retaining a larger subset of the full ABCD sample (N=9,753), replicated the effect sizes across all brain-wide associations (see Methods).

In Fig. 1a,b, we show the univariate associations between cortical thickness/RSFC and two extensively studied behavioral phenotypes, cognitive ability (NIH Toolbox total composite score) and psychopathology (Child Behavior Checklist^24^ [CBCL] total problem score; see Methods & Extended Data Table S1). In the full denoised sample (N=3,928), across all brain-wide associations, the median effect size was |r|=0.01 (Extended Data Fig. S1). The 99^th^ percentile (largest 1%) of all possible brain-wide associations was |r|>0.06. The strongest correlation between a brain metric and a behavioral measure was |r|=0.16 (edge-level: RSFC with cognitive ability). Across all brain-wide associations, the single largest correlation that replicated out-of-sample was |r|=0.14 (edge-level: RSFC with crystallized intelligence composite score [NIH Toolbox]^25^). For exemplar behavioral phenotypes (cognitive ability, psychopathology), the strongest associations were distributed across sensorimotor and association cortex (Fig. 1c,d). These patterns of brain-wide associations suggest that cognitive ability and psychopathology are supported by widely distributed circuitry.

**Fig 1.**
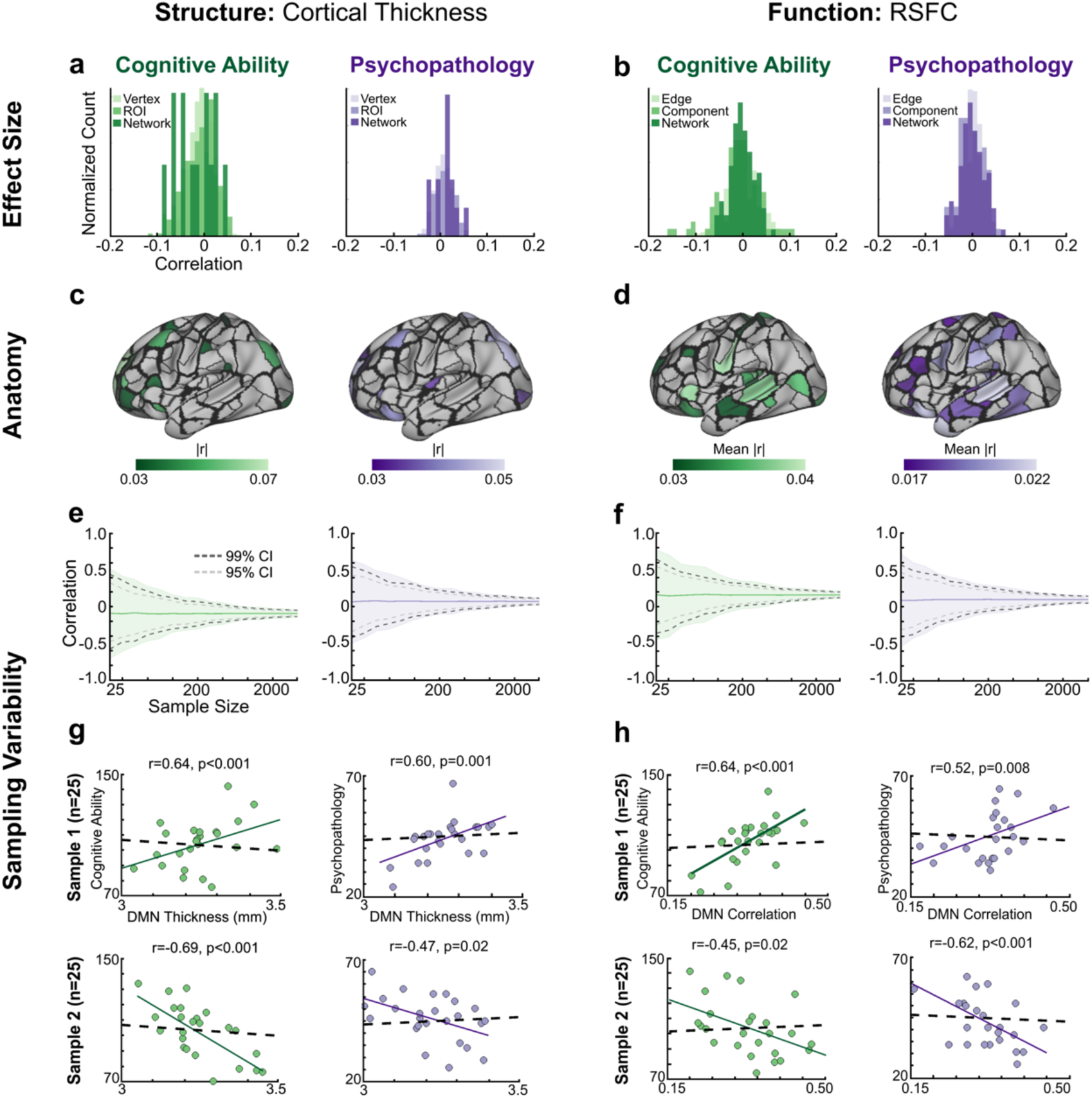
Effect Sizes and Sampling Variability of Univariate Brain-Wide Associations as a Function of Sample Size. Correlations of **(a)** cortical thickness with cognitive ability (left; green), psychopathology (right; purple) at different levels of analysis (vertex, ROI, network) and **(b)** RSFC with cognitive ability (left; green), psychopathology (right; purple) at different levels of analysis (edge, network, component). Brain regions with the largest brain-wide associations (top 10%) for **(c)** cortical thickness with cognitive ability (left; green), psychopathology (right; purple); **(d)** RSFC with cognitive ability (left; green), psychopathology (right; purple). Sampling variability (range of observed correlations; 1,000 resamplings per sample size) for the largest brain-wide associations; **(e)** cortical thickness with cognitive ability (left; green), psychopathology (right; purple); **(f)** RSFC with cognitive ability (left; green), psychopathology (right; purple). Solid lines represent the mean across 1,000 simulated studies (resamplings). Shading represents the minimum and maximum correlation across the 1,000 resamples for a given sample size. Gray dotted line denotes the 95% confidence interval (CI). Black dotted line denotes the 99% CI. **(g)** Default Mode network (DMN) cortical thickness with cognitive ability (left; green), psychopathology (right; purple) correlations from two N=25 samples, in which significant, but inaccurate correlations with opposite signs are observed (top *vs.* bottom). Black dotted line denotes least squares fit from the full sample **(h)** Default Mode network (DMN) RSFC with cognitive ability (left; green), psychopathology (right; purple) correlations from two N=25 samples, in which significant, but inaccurate correlations with opposite signs are observed (top *vs.* bottom).

Smaller brain-wide association studies have reported larger correlations than the largest effect size in the ABCD Study^26^. How can that be? In Fig. 1e,f, we show that sampling variability (99% confidence interval [CI] of observed correlations) alone generates nominally significant (p<0.05), but inflated correlations, which would then be falsely reported^2,27^. We charted sampling variability as a function of sample size (N=25 to 3,928) for the strongest brain-wide associations as defined in the full sample (N=3,928, strict denoising). We chose the strongest association (RSFC with cognitive ability: r=0.16 in full sample, r=0.14 out-of-sample) as an illustrative example. At the median sample size of existing BWAS (N=25), the 99% CI for RSFC and cognitive ability associations ranged from r=-0.39 to r=0.62 (Fig. 1e,1f), demonstrating how sampling variability alone can account for a broad range of observed brain-wide associations. Although sampling variability decreases with increasing sample size, a sample as large as N=2,000 still contains measurable sampling variability. For example, the sampling variability (99% CI) of the strongest RSFC, cognitive ability association (r=0.16) at N=2,000 was 0.10-0.22.

In Fig 1g,h, we provide examples of the potentially deleterious effects of sampling variability using the heavily-studied association between default mode network (DMN) and cognitive ability/psychopathology. Here, two N=25 subsamples (e.g., cortical thickness with cognitive ability) can reach the *opposite* conclusion (r=0.64 vs. r=-0.69; full sample: r=0.08) with nominal statistical significance (both p<0.001), solely due to sampling variability of the association. The effects of sampling variability shown in Fig. 1e,f generalized to all brain-wide associations (Extended Data Fig. S2,S3) as well as behavior-behavior correlations (e.g. cognitive ability and psychopathology correlation range = 1.25 at N=25, Extended Data Fig. S4).

### Effect Sizes and Sampling Variability Replicate in Single Site Datasets

Given the pediatric (9-10 yrs) and multi-site sample of the ABCD Study, we sought to replicate effect size estimates in a single site, single scanner, adult dataset (HCP: N=1,200; N=877 post-denoising; age-range: 22-35 years). In HCP data, we identified modestly larger correlations between RSFC and NIH Toolbox subscales (max. r=0.20) than those observed in the ABCD data (max. r=0.16, full denoised sample). However, subsampling ABCD data to match the HCP sample size (N=877) inflated the ABCD’s maximum correlation from 0.16 to 0.20 and generated a nearly identical effect size distribution as for the HCP Study (Extended Data Fig. S5a), indicating that brain-wide associations in the 21-site pediatric ABCD study are equivalent in magnitude to those in the single site/scanner, adult HCP study. These results suggest that any effect sizes >r~0.16 in HCP data may still be inflated due to sampling variability. In fact, even across 100 bootstrapped split-half ABCD samples (N=1,964 in each half), the top 1% largest brain-behavioral phenotype correlations were inflated by r=0.043 (r=0.027 in N=4,303 split halves, no exclusion for data quality), on average (Extended Data Fig S5b). Although measurable sampling variability remains, sample sizes of N⪆2,000 (post-denoising) can provide more stable estimates of the associations between brain and behavioral measures.

To ameliorate any concerns that sampling variability may be inflated by the multi-site nature of ABCD, we directly compared sampling variability between the HCP and ABCD datasets (Extended Data Fig. S6a), and between a single ABCD site (N=603) and the remaining sites (Extended Data Fig. S6b). In both cases, sampling variability was equivalent between single site and multi site samples, providing evidence that sampling variability was robust to site effects.

### Statistical Errors Are Ubiquitous in Typically Sized Brain-Wide Association Studies (BWAS)

In Fig. 2, we show that the combination of observed BWAS effect sizes and sampling variability generates statistical errors. Specifically, we plotted statistical error rates (false negative [type 2 error], false positive, [type 1 error], correlation magnitude inflation [type M], and sign errors [type S])^28^ as a function of sample size for all univariate brain-wide associations (Extended Data Fig. S7 displays each MRI measure separately). For typical BWAS samples (e.g., N≤200), the false negative rate was at least 76% at p<0.05 (Fig. 2a) with a concurrent false positive rate of 6.1% (Fig. 2b), meaning typically-sized samples mischaracterize a significant correlation as a non-significant correlation at least 76% of the time in an uncorrected, hypothesis-driven analysis. Thus, at p<0.05, N≤200 samples only achieved a maximum power of 24% (power= 1-false negative rate). Sample sizes of at least 2,200 subjects were required to be 80% powered to detect the largest 1% of univariate brain-wide associations (r=0.06) at *p*<0.05 (Extended Data Fig. S8a).

**Fig 2.**
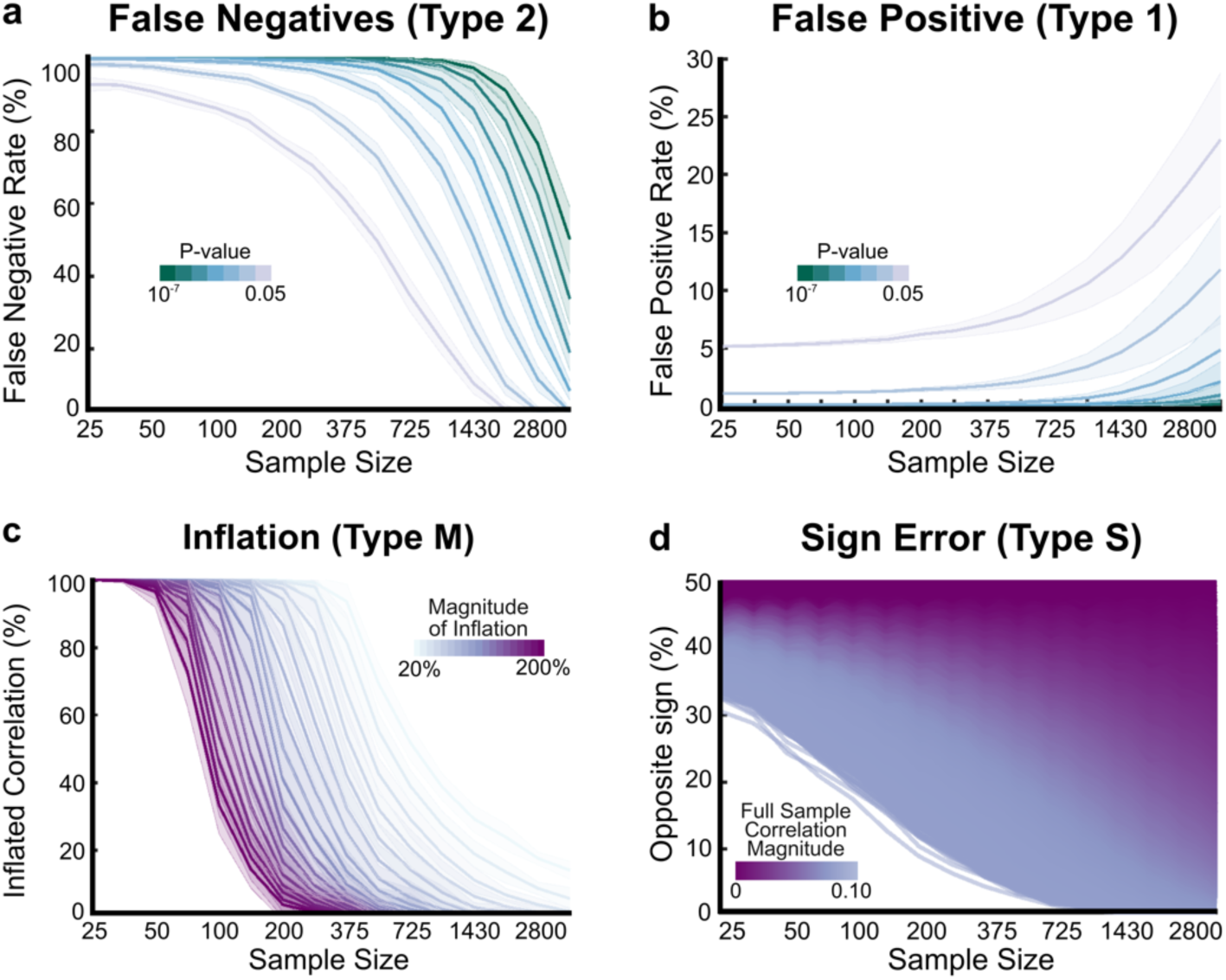
Statistical Errors in Brain-Wide Associations as a Function of Sample Size. **(a)** False negative rate in RSFC (edgewise) correlations with cognitive ability varying as a function of p-value. The most stringent p-value (10^−7^) is equivalent to a Bonferroni correction across all 77,421 RSFC pairs, whereas p<0.05 is equivalent to no multiple comparisons correction. **(b)** False positive rate in RSFC correlations with cognitive ability varying as a function of p-value. **(c)** Inflation rate of the RSFC correlation with cognitive ability, depicting the probability of observing an inflated correlation as a function of sample size. Color gradient represents the magnitude of inflation. **(d)** Probability of observing the opposite sign of the correlation observed between RSFC and cognitive ability in the full sample (N=3,928) as a function of sample size. Color gradient represents the effect size of the correlation in the full sample.

When correcting for multiple comparisons (Bonferroni), 9,500 subjects were required to be 80% powered to detect the largest 1% (r=0.06) of correlations (Extended Data Fig S8b). If an N≤200 sample correctly rejected the null hypothesis (at most 24% at p<0.05; 0.1% at p<10^−7^), there was a 50% chance that the magnitude of the correlation was inflated by at least 100% (Fig. 2c). Additionally, across all brain-wide associations, samples with an N≤200 had at least a 40% chance of reporting correlations with the wrong sign (Fig. 2d), meaning that they reach the opposite conclusion 40% of the time.

To further test the reproducibility of univariate brain-phenotype associations, we determined the out-of-sample replication of brain-wide associations using established discovery (N=1,964) and replication (N=1,964) ABCD sets, matched across a broad range of demographic factors^29^ (Extended data Fig. S9). In N≤200 samples, univariate models never replicated out-of-sample (R^2^ range: −0.018 - 0.005). In the full replication sample (N=1,964), the top 0.5% strongest brain-wide associations achieved modest out-of-sample replication, explaining 1% of the variance in out-of-sample data (R^2^=0.01).

### The “Underpowered Correlation Paradox”

Paradoxically, at small sample sizes, the largest correlations are the most erroneous, yet the most likely to be significant, and therefore the most likely to be published. This correlation paradox occurs because typical BWAS samples (e.g., N=200) are only sufficiently powered to detect statistical significance for correlations that are larger than the largest correlation (r=0.16) observed in the full ABCD study sample. Thus, typically sized studies (e.g., N=200) that report significant brain-wide associations necessarily must have inflated effect sizes (r>0.16). This happens by chance, as sampling variability enables the detection of a significant (p<0.05) yet inflated correlation (Fig 1e,f). When attempting to replicate a significant inflated correlation - due to the central tendency of the effect size distribution across studies - the most likely outcome would be to observe a correlation of r~0.10 (value in full sample), thus failing to replicate the original inflated effect. Publication bias (only publishing significant effects) and reliance on null hypothesis-testing for inference would likely prevent the replication result from being published, perpetuating inflated effect sizes that form the basis for subsequent power- and meta-analyses.

### Multivariate Associations Can Improve Reproducibility in Large Samples

Multivariate approaches (SVR [Support Vector Regression], CCA [Canonical Correlation Analysis]) are thought to be more powerful for discovering brain-wide associations (Fig. 3)^30^. SVR detects multivariate brain patterns (as opposed to single brain features in univariate approaches) that predict a *single* behavioral phenotype (e.g., cognitive ability), while CCA enables the discovery of weighted multivariate brain patterns that predict weighted combinations of *multiple* behavioral phenotypes (e.g., all NIH Toolbox subscales). To examine multivariate brain-wide associations as a function of sample size, we fit SVR and CCA models on subsamples of the discovery set (in-sample) and subsequently tested their out-of-sample replicability on the full replication set. Replicability was quantified as the correlation (r) between observed behavioral scores in the replication dataset and predicted behavioral scores from models fit on the discovery set.

**Fig 3.**
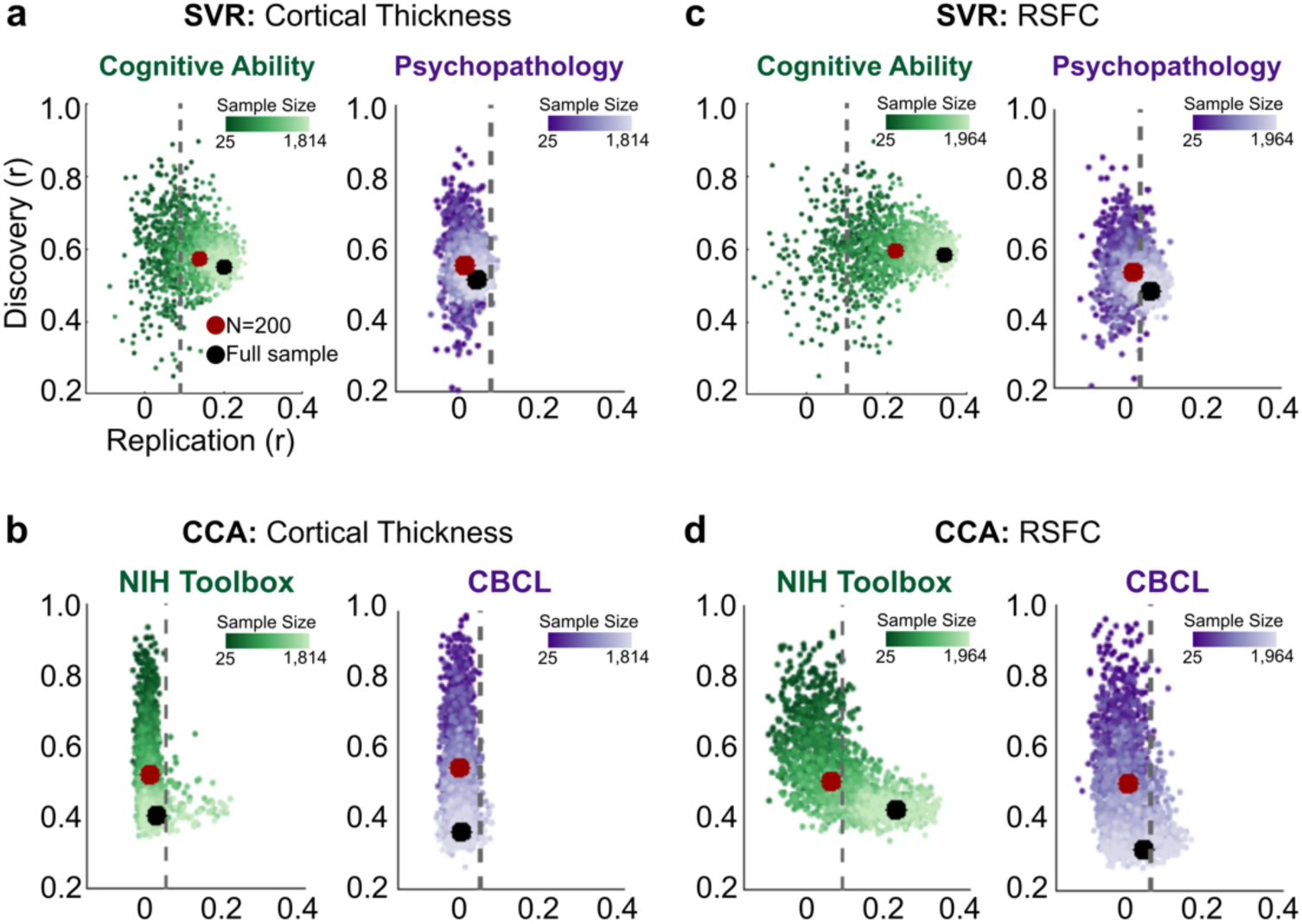
Multivariate Brain-Wide Associations. Discovery (in-sample) multivariate (SVR, CCA) correlations (y-axes) as a function of out-of-sample replication (x-axes) and sample size (dark-to-light color gradient) for **(a)** the multivariate (SVR) associations of cortical thickness with cognitive ability (green; left), psychopathology (purple; right). **(b)** The multivariate (CCA) association of cortical thickness with the NIH Toolbox (green; left), CBCL (purple; right). **(c)** The multivariate (SVR) relationship of RSFC with cognitive ability (green; left), psychopathology (purple; right). **(d)** The multivariate (CCA) relationship of RSFC and the NIH Toolbox (green; left), with CBCL (purple; right). Gray dotted line represents the threshold for significant replication (p<0.01), determined through permutation testing on the full sample for each brain-wide association. Circles indicate mean replication (r) across 100 bootstrap samples (with replacement) at an N=200 (red dot) and the full sample (black dot). Full sample sizes: cortical thickness N=1,814; RSFC N=1,964.

As shown in Fig 3a,b, multivariate models fit on cortical thickness data in typical BWAS samples only replicated when using SVR to predict cognitive ability (r=0.14, null r=0.09; see Extended Data Fig. S10, S11 for SVR feature and hyperparameter selection). Other cortical thickness multivariate models never replicated with typical BWAS samples (N=200), despite large brain-phenotype associations in the discovery sample (e.g., CCA range at N=200 for discovery r=0.37-0.81; Fig. 3b). However, as sample sizes approached consortium-levels (i.e., the full discovery sample N=1,814), SVR associations between cortical thickness and cognitive ability achieved significant replication (observed r=0.20, 99% confidence of the null r=0.09, *p*<0.01; Fig. 3a). This was the largest observed replication between cortical thickness and any behavioral phenotype. Using SVR, replication significance was not achieved for psychopathology (r=0.05, null *r*=0.08, *p*>0.01; Fig. 3a) or for cortical thickness associations with the NIH Toolbox or the CBCL (Fig. 3b).

Consistent with the requirement for consortium-level sample sizes for reproducibility, multivariate models fit on RSFC data in typical BWAS samples mostly resulted in replication failures. For example, SVR models fit on RSFC data from N=200 subsamples of the discovery set (Fig. 3c) replicated with cognitive ability (RSFC/cognitive ability out-of-sample r=0.15, null r=0.08), but failed to replicate with psychopathology (RSFC with psychopathology r=0.05, null r=0.06; Fig. 3c). At N=200, CCA models never replicated (all out-of-sample r<0.07; Fig 3d), despite achieving high correlations in the discovery sample (RSFC with NIH Toolbox r=0.35-0.66; RSFC with CBCL r=0.32-0.66; Fig. 3d). As sample sizes approached consortium-levels (N=1,964), significant replication between RSFC and cognitive ability with SVR was observed (r=0.34, null r=0.10; Fig. 3c). The SVR association between RSFC and cognitive ability was the largest replicated brain-wide effect across all measures and is consistent with other recent large-scale prediction efforts^31,32^. Similarly, CCA associations between RSFC and the NIH Toolbox fit on the full discovery sample achieved significant out-of-sample replication (r=0.22, null r=0.09, *p*<0.01; Fig. 3d), but not between RSFC and CBCL (r=0.06, null r=0.08 *p*>0.01; Fig. 3d).

Multivariate out-of-sample replication (max. r=0.34) was superior to univariate out-of-sample replication at consortium-level sample sizes (N~2,000), but was still low-to-moderate for most behavioral phenotypes. Out-of-sample replication was generally maximized by using a relatively low-dimensional feature space (e.g., principal components accounting for 20-50% of cumulative variance; Extended Data Fig. S12, S13), reaffirming that brain-wide associations are represented in widely-distributed circuitry^33^, consistent with the univariate results (Fig 1c,d). Moreover, multivariate replicability was robustly linked to univariate effect sizes, such that multivariate replicability and univariate effect sizes were strongly correlated (r=0.79, p<0.001, Extended Data Fig. S14).

### Towards Reproducible Brain-Wide Association Studies through Large-Sample Consortia

Human neuroscience’s goal of reproducibly linking the brain to behavioral phenotypes requires sample sizes only recently achieved by consortia (e.g., ABCD, N>10,000). Typical sample sizes collected by single investigators have been drastically underpowered for brain-wide association studies (BWAS), thus largely failing to replicate.

Similar to even larger genomics consortia, highly collaborative and strategic human neuroimaging studies such as the ABCD, HCP, The All of Us Research Program^34^, and the UK Biobank are empowering neuroscientists to identify robust and reproducible brain-wide associations between brain and behavioral measures (Fig. 4). As in other population-based disciplines, many of which have substantially smaller effect sizes (e.g., individual gene variants: <0.01% variance explained), consortium efforts are promoting reproducibility, methods harmonization,^35^ and the collection of sample sizes with greater statistical power. For example, genomics researchers realized that a handful of well-characterized genes were likely false discoveries resulting from underpowered studies^21^. Large consortia efforts in genomics have led to sample sizes exceeding 1 million individuals^36^ with standardized methodological pipelines^37^. These large discovery sets are used to train whole-genome models that are subsequently used to answer more detailed questions in independent samples^18^. Currently, brain-wide association studies are trapped in the initial discovery phase. Larger samples will inevitably allow multivariate models, both those currently in use and future improvements currently being developed^38–41^, to be employed by individual research groups.

**Fig 4.**
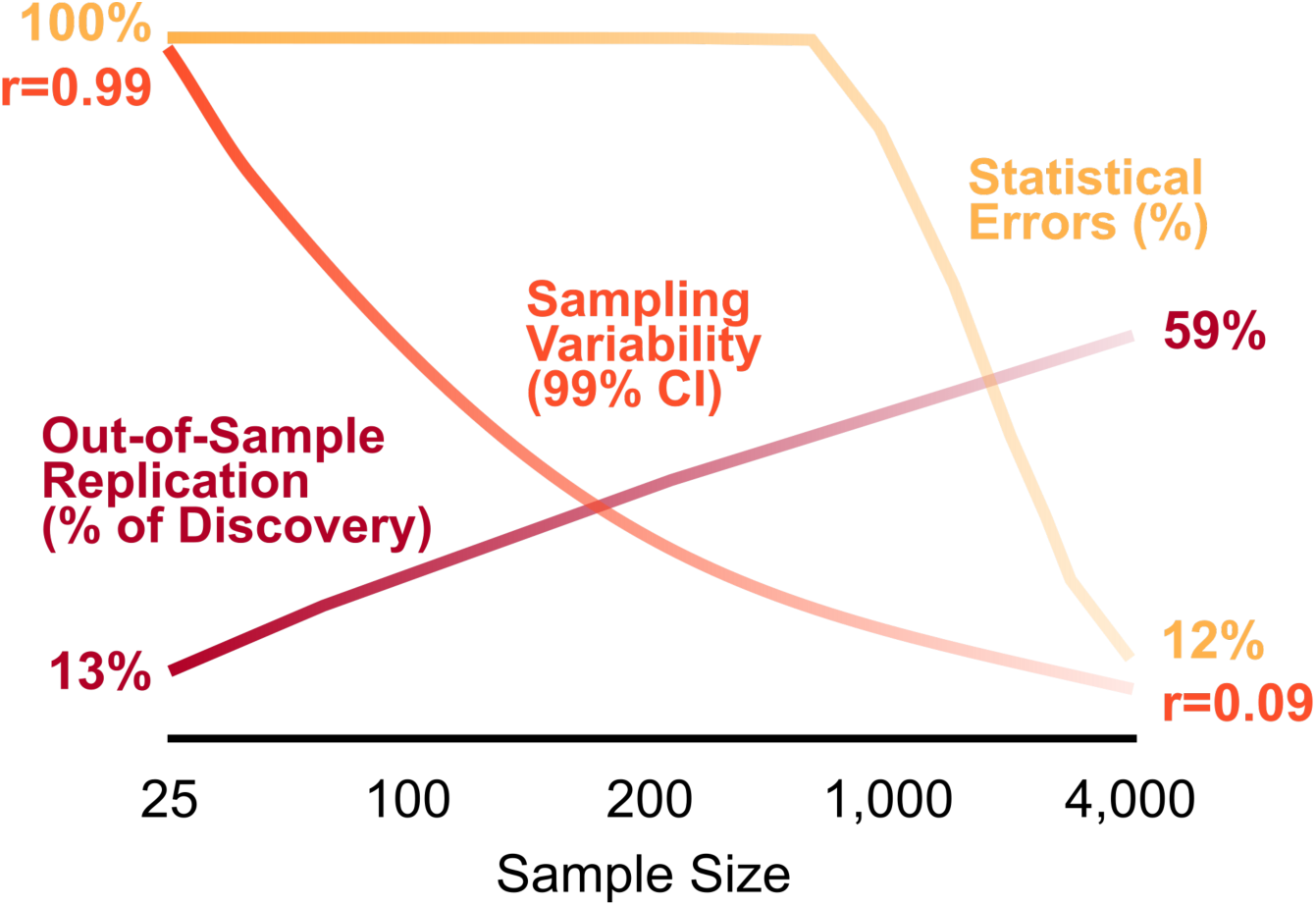
Influence of Sample Size on the Robustness of Brain-Wide Associations (BWAS). Trajectories of sampling variability (99% confidence interval; orange), statistical error rates (cumulative sum of false negatives, false positives, effect size inflation, sign errors; yellow), and SVR out-of-sample replication (percentage of the magnitude of the full sample discovery [in-sample] correlation [% Pearson r]; dark red) as a function of sample size. Sample size (e.g 4,000) represents a full sample (discovery + replication datasets of 2,000 each).

Smaller, investigator-initiated neuroimaging studies will continue to be just as important for human neuroscience as they were in the days when samples of N>10,000 for brainwide-wide association studies (BWAS) were still an impossibility. Larger sample consortium studies of brain-wide associations were made possible by a myriad of independent studies that created the technological, analytical, and theoretical frameworks adopted by the ABCD consortium. Smaller neuroimaging studies have provided blueprints for reducing MRI artifacts^42^, increasing usable data^43^, and boosting reliability within individuals ^44,45^. Indeed, the amount of RSFC data collected per subject in ABCD was based on an N=1 study^46^. There is no one-size-fits-all sample size for neuroimaging studies^47^; instead the appropriate sample size must always be determined by the study goals. Just as one should not use a N>10,000 consortium to experiment with novel ideas, technologies, or methods, one should not conduct brain-wide association studies - BWAS - with anything but the very largest, currently available samples. Studying brain-wide associations in smaller samples requires well-controlled, hypothesis-driven experimental manipulations (e.g., interventions, lesions) to boost effect sizes^48^. Trailblazing smaller studies should always seek replication in substantially larger sample sizes^49^, and contribute study data to repositories for future data aggregation.

Ultimately, making brain imaging measures maximally relevant to human behavioral phenotypes requires ever larger population-based samples, as showcased by genomics^18^. By building large multi-site consortia on the wisdom gleaned from previous smaller studies, we are poised to take the long-awaited leap towards robustly and reproducibly linking the brain’s structure and function to the complex behavioral phenotypes that shape our lives.

## Methods

### Sample

This project utilized the ABCD-BIDS processed dataset consisting of RSFC data from 10,259 participants through the ABCD-BIDS Community Collection (ABCD collection 3165; https://github.com/ABCD-STUDY/nda-abcd-collection-3165) and demographic and behavioral data from 11,572 9-10 year old participants from the ABCD 2.0 release^3^. We also utilized the ABCD reproducible matched samples (ARMS), available in ABCD collection 3165, that divided individuals from the full behavioral sample (N=11,572) into discovery (N=5,786) and replication (N=5,786) sets, which were matched across 9 variables: site location, age, sex, ethnicity, grade, highest level of parental education, handedness, combined family income, and prior exposure to anesthesia. Family members (e.g., sibling pairs, twins, and triplets) were kept together in the same set and the two sets were matched to include equal numbers of single participants and family members. These split datasets were used for replicability analyses.

Head motion can systematically bias neuroimaging studies^50^. However, these systematic biases can be addressed through rigorous head motion correction^51^. Therefore, we used strict inclusion criteria with regard to head motion. Specifically, inclusion criteria for the current project (see ref^52^ for broader ABCD inclusion criteria) consisted of at least 600 frames (8 minutes) of low-motion (filtered FD<0.08) RSFC data. Our final dataset consisted of RSFC data from a total of N=3,928 youth across the discovery (N=1,964) and replication (N=1,964) sets. The final discovery and replication sets did not differ in mean FD (∆*M*=0.002, *t*=0.60, *p*=0.55) or total frames included (∆M=6.4, *t* =0.94, p=0.35). The subject lists for ARMS samples and our associated matrices will be released in the ABCD-BIDS Community Collection (ABCD collection 3165) for community use.

### MRI acquisition

Imaging was performed at one of 21 sites within the United States, harmonized across Siemens Prisma, Philips, and GE 3T scanners. Details on image acquisition can be found in^52^. Twenty minutes (4 × 5 minute runs) of eyes-open (passive crosshair viewing) resting state data were acquired to ensure at least 8 minutes of low-motion data. All resting state scans fMRI scans used a gradient-echo EPI sequence (TR = 800 ms, TE = 30 ms, flip angle = 90°, voxel size = 2.4 mm^3^, 60 slices). Head motion was monitored using Framewise Integrated Real-time MRI Monitor (FIRMM) software at many of the Siemens sites^43^.

### ABCD-BIDS (HCP-style CIFTI) processing overview

All processing was completed with the newly released and freely available ABCD-BIDS pipelines (https://github.com/DCAN-Labs/abcd-hcp-pipelines). The ABCD-BIDS pipelines are modified from the original HCP pipelines^53^. Briefly, this pipeline comprises six stages. 1) PreFreesurfer normalizes anatomical data. This normalization entails brain extraction, denoising, and then bias field correction on anatomical T1 and/or T2 weighted data. The ABCD-HCP pipeline includes two additional modifications to improve output image quality. ANTs^54^ DenoiseImage models scanner noise as a Rician distribution and attempts to remove such noise from the T1 and T2 anatomical images. Additionally, ANTs N4BiasFieldCorrection attempts to smooth relative image histograms in different parts of the brain and improves bias field correction. 2) FreeSurfer^7^ constructs cortical surfaces from the normalized anatomical data. This stage performs anatomical segmentation, white/grey and grey/CSF cortical surface construction, and surface registration to a standard surface template. Surfaces are refined using the T2 weighted anatomical data. Mid-thickness surfaces, which represent the average of white/grey and grey/CSF surfaces, are generated here. 3) PostFreesurfer converts prior outputs into an HCP-compatible format (i.e. CIFTIS) and transforms the volumes to a standard volume template space using ANTs nonlinear registration, and the surfaces to the standard surface space via spherical registration. 4) The “Vol” stage corrects for functional distortions via reverse-phase encoding spin-echo images. All resting state runs underwent intensity normalization to a whole brain mode value of 1000, within run correction for head movement, and functional data registration to the standard template. Atlas transformation was computed by registering the mean intensity image from each BOLD session to the high resolution T1 image, and then applying the anatomical registration to the BOLD image. This atlas transformation, mean field distortion correction, and resampling to 3-mm isotropic atlas space were combined into a single interpolation using FSL’s^55^ applywarp tool^56^. 5) The “Surf” stage projects the normalized functional data onto the template surfaces which is described below. 6) We have added an fMRI and fcMRI preprocessing stage, “DCANBOLDproc” which is also described below. 7) Last, an Executive Summary is provided for easy subject level QC across all processed data.

### fMRI Surface (Surf) processing

The BOLD fMRI volumetric data were sampled to each participant’s original mid-thickness left and right-hemisphere surfaces constrained by the grey-matter ribbon as described in ^53^. Once sampled to the surface, time courses were deformed and resampled from the individual’s original surface to the 32k fs_LR surface in a single step. This resampling allows point-to-point comparison between each individual registered to this surface space. These surfaces were then combined with volumetric subcortical and cerebellar data into the CIFTI format using Connectome Workbench ^57^, creating full brain time courses excluding non-gray matter tissue. Finally, the resting-state time courses were smoothed with 2mm full-width-half-maximum (FWHM) kernel applied to geodesic distances on surface data and euclidean distances on volumetric data.

### (DCANBOLDproc) preprocessing

Additional BOLD preprocessing steps were executed to reduce spurious variance unlikely to reflect neuronal activity^42^. First, a respiratory filter was used to improve FD estimates calculated in the volume (“vol”) stage. Second, temporal masks were created to flag motion-contaminated frames using the improved FD estimates^50^. Frames with a filtered FD>0.3mm were flagged as motion-contaminated for nuisance regression only. After computing the temporal masks for high motion frame censoring, the data were processed with the following steps: (i) demeaning and detrending, (ii) interpolation across censored frames using least squares spectral estimation of the values at censored frames^51^ so that continuous data can be (iii) denoised via a GLM with whole brain, ventricular, and white matter signal regresssors, as well as their derivatives. Denoised data were then passed through (iv) a band-pass filter (0.008 Hz<f<0.10 Hz) without re-introducing nuisance signals^58^ or contaminating frames near high motion frames.

### Generation of Resting State Functional Connectivity (RSFC) matrices

For each subject, high motion frames (filtered FD>0.08) were censored. The timeseries of BOLD activity for each region of interest (333 cortical ROIs from Gordon et al.^59^; 61 subcortical ROIs from Seitzman et al.^60^) was correlated to that of every other ROI, forming a 394 × 394 correlation matrix, which was subsequently Fisher z-transformed. For network level analyses, correlations were averaged across previously defined canonical functional networks^59^. Inter-individual difference connectome-wide spatial components, which are not bound by network boundaries^29,33^, were computed by performing principal component analysis (PCA) on a matrix composed of all ROI × ROI pairs (edges) from each participant.

### Generation of Cortical Thickness Metrics

For each subject, cortical thickness was extracted from 59,412 cortical vertices. For ROI level matrices, cortical thickness was averaged within each cortical parcel (N=333)^59^. For network level matrices, cortical thickness was averaged within each cortical network (N=13)^59^. Inter-individual spatial components were computed by performing principal component analysis on a matrix composed of all cortical vertices from each participant.

### Behavioral and Demographic Data

The ABCD study population is well-characterized with hundreds of demographic, physical, cognitive, and mental health variables^61^. The current project examined the associations between 41 of these variables (Extended Data Table S1) and brain structure (cortical thickness) and function (RSFC). Behavioral and demographic variables were selected to reflect the primary domains of interest, cognition (individual measure and composite scores from the NIH Toolbox)) and mental health (syndrome scales and broad-band psychopathology factors from the CBCL), as well as demographic/physical variables relevant to development (e.g., age) and health (e.g., BMI).

### Behavioral and Demographic Covariates

The primary goal of this project was to study how the pairing of brain-behavioral phenotype effect sizes and sampling variability (random variation across samples, as opposed to systematic variation threatening causal inference^62^) can account for wide-spread replication failures. As a result, our results focus on simple bivariate associations (correlation) and canonical multivariate models linking brain structure and function to behavioral and demographic variables without covariate adjustment. However, we note that the ABCD subsamples (ARMS; see Above) we used for replication analyses are matched for salient demographic factors (site location, age, sex, ethnicity, grade, highest level of parental education, handedness, combined family income, and prior exposure to anesthesia; see above). Also, where possible, ABCD-distributed age-corrected scores were used, given 1) well-established age-related changes in these measures and 2) age-corrected scores improved normality for many measures (e.g., CBCL syndrome scales and broadband factors) symptom presentation).

### Capture of Behavioral and Demographic Data

The ABCD Data Analysis and Informatics Center (DAIC) has released an online tool called DEAP (Data Exploration and Analysis Portal), which can be accessed at https://deap.nimhda.org/. In this manuscript, we introduce an additional tool called ABCDE, ABCD Boolean Capture Data Explorer, which we have used for preparation of the data herein. ABCDE complements DEAP by allowing for finer-grained control of data extraction on the researcher’s own computer rather than through a web portal. The source code and documentation can be accessed at https://gitlab.com/DosenbachGreene/abcde.

### Univariate Brain-Behavioral Phenotype Correlations

For each brain measure at a given level of organization, we correlated the brain measures (structure: cortical thickness; function: RSFC) with each behavioral variable. Cognitive ability (total composite score on the NIH Toolbox) and psychopathology (total score on the Child Behavior Checklist [CBCL]) are presented in the main text; all others are in the supplemental material. Correlations between brain and behavior were generated for RSFC at the *edge level* (ROI-ROI pair [N=77,421]), *network level* (average of RSFC within/between each network [n=105]), and *component level* (principle component weights [N=100]). To extract individual-specific components, we vectorized each participant’s RSFC matrix, concatenated the vectorized matrices, and submitted them to singular value decomposition (Matlab’s svd.m function). Correlations between brain and behavioral phenotypes were generated for cortical thickness at the *vertex level* (N=59,412), *ROI level* (N=333), and *network level* (N=13). Repeat analyses employing less rigorous motion censoring, and thus retaining a larger subset of the full ABCD sample (N=9,753), replicated the effect sizes (maximum r=0.16, 99^th^ percentile [largest 1%] of effect sizes: |r|>0.06).

### Resampling Procedures

To examine the distribution of correlations for iteratively larger sample sizes, we randomly selected subjects with replacement from the full sample (N=3,928, post denoising) at logarithmically spaced sample sizes (16 intervals: N=25, 33, 50, 70, 100, 135, 200, 265, 375, 525, 725, 1,000, 1,430, 2,000, 2,800, 3,928). For cortical thickness data, the full sample contained (N=3,604) in the same sampling bins, with the exception of the final bin (full sample), which contained 3,604 participants. At each sample size, we randomly sampled subjects 1,000 times, resulting in 16,000 brain-behavioral phenotype resamplings for each brain-behavioral phenotype correlation. For multivariate approaches, 100 bootstrap samples were computed across the same logarithmically-spaced sample sizes. We note that the iterations were reduced for multivariate methods due to their high computational costs. In addition, the analyses are focused on mean estimates, not the full distribution. For highlighting the effects of sampling variability (Fig. 1e,f), we extracted the brain-behavioral phenotype correlation with the largest effect size for each imaging measure (cortical thickness, RSFC) and exemplar behavioral phenotypes (cognitive ability, psychopathology). The sampling variability (range of possible correlations, 99% confidence interval and 95& confidence interval) at each sampling interval for correlations between RSFC and cortical thickness with cognitive ability and psychopathology are presented in the main text (Fig. 1e,f); correlations between brain measures and other behaviors can be found in the Extended Data (Fig. S2,S3).

### Empirical Quantification of Sampling Variability: Example Studies at N=25

Using the outputs from the resampling procedures above, we used the 1,000 resamplings with N=25 to examine the correlation between the default mode network (DMN) and cognitive ability (total composite score on the NIH Toolbox), as well as the default mode network (DMN) and psychopathology (total problem score on the CBCL), for both cortical thickness and RSFC. To demonstrate how sampling variability affects correlations, the 1,000 resamples were ranked by effect size. Subsequently, two samples were chosen from the top 10 studies (in terms of effect size); one with a significant positive association (Sample 1) and one with a significant negative association (Sample 2).

### Behavior-Behavior Correlations

To examine the range of sampling variability as a function of sample size between 41 behavioral and demographic measures, we randomly selected subjects with replacement from the full behavioral sample (N=11,572) at logarithmically spaced sample sizes (9 intervals: N=25, 50, 100, 200, 500, 1,000, 2,000, 4,000, 9,000). At each interval, we randomly sampled subjects 1,000 times, resulting in 9,000 phenotype-phenotype correlation resamplings for each association. For each association, we quantified sampling variability at each sampling bin as the range of correlations observed through the resampling procedure.

### False Positives and False Negatives

False negative (Fig. 2a) and false positive (Fig. 2b) rates were derived through a resampling procedure (see <u>Resampling Procedures</u>) for all edge-level brain-wide associations. For each sample size bin (16 total), we randomly sampled “N” individuals (1,000 subsamples) and computed the brain-behavioral phenotype correlation and associated p-value. A correlation was deemed “significant” if it survived a threshold of p<0.05, Bonferroni corrected (p<10^−7^; corrected across 77,421 ROI-ROI pairs) in the full sample (cortical thickness N=3,604, RSFC N=3,928). At each sample size, if a correlation in the full sample was *not* significant, we determined the percentage of studies that resulted in a “significant” correlation (false positive) across a broad range of p-values (10^−7^ to 0.05). Conversely, if a correlation in the full sample *was* significant (p<10^−7^), we determined the percentage of studies that resulted in a “non significant” correlation (false negative) across a broad range of p-values (10^−7^ to 0.05).

### Correlation Inflation

For each univariate brain-wide association in the full sample (N=3,928) at the edge level, we determined whether or not a correlation was significant (p<10^−7^, Bonferonni corrected for multiple comparisons [p=0.05/77,421]). For each significant correlation, across the 1,000 subsamples within a sampling bin, we extracted the percentage (out of 1,000) of significant correlations (p<0.05) that were inflated across a range of inflation thresholds (20-200%, in steps of 1%).

### Correlation Sign Errors

Each brain-wide association was extracted from the full sample as a reference. Across the 1,000 subsamples within a sampling bin, we determined the percentage of correlations that had the opposite correlation sign as the correlation sign in the full sample.

### Human Connectome Project (HCP) Replication: Effect Sizes

We used data from N=877 individuals from the Human Connectome Project (HCP) 1,200 Subject Data Release (aged 22-35 years). A custom Siemens SKYRA 3.0T MRI scanner and a custom 32-channel Head Matrix Coil were used to obtain high-resolution T1-weighted (MP-RAGE, 2.4 s TR, 0.7×0.7 × 0.7 mm voxels) and BOLD contrast sensitive (gradient-echo EPI, multiband factor 8, 0.72 s TR, 2×2×2mm voxels) images from each subject. The HCP used sequences with left-to-right (LR) and right-to-left (RL) phase encoding, with a single RL and LR run on each day for two consecutive days for a total of four runs^63^. MRI data were preprocessed as previously described^60^. All HCP data are available at https://db.humanconnectome.org/. Similar to the ABCD data, we extracted the timeseries from a total of 394 cortical and subcortical ROIs, correlated and Fisher z-transformed them. Behavioral data from the NIH Toolbox were correlated with each edge of the correlation matrix across subjects. Across all NIH Toolbox subscales, the tails of the distributions of the resulting brain-phenotype correlations were compared to 100 subsampled ABCD brain-behavioral phenotype correlations (N=877, matching HCP sample size). In Extended Data Fig. S5a, we show the distributions of brain-behavioral phenotype correlations for ABCD and HCP data for each NIH Toolbox subscale.

### Human Connectome Project Replication: Sampling Variability

To quantify the degree of sampling variability in single site, single scanner HCP data compared to multi-site, multi-scanner ABCD data, we subsampled ABCD RSFC data to match HCP sample sizes (N=877). For each dataset, we ran the resampling procedure detailed in “Resampling Procedures” above (12 intervals: N=25, 33, 50, 70, 100, 135, 200, 265, 375, 525, 725, 875) across all NIH Toolbox subscales. The range of correlations and 95% confidence interval observable in each sampling bin are shown in Extended Data Fig. S6a for both HCP and ABCD data.

### Single-site ABCD vs. Multi-site ABCD: Sampling Variability

We directly compared single site ABCD data (site 16; N=603) with multi-site ABCD data (N=3,325, 20 sites, site 16 was excluded) using the same resampling procedure (10 intervals: N=25,33, 50, 70, 100, 135, 200, 265, 375, 525) as before on the associations between RSFC and all NIH Toolbox subscale (Extended Data Fig. S6b). The range of correlations and 95% confidence intervals in each resampling bin is shown in Extended Data Fig. S6b for single-site and multi-site ABCD data.

### Replication of Univariate Associations

Univariate replication was performed across primary analysis variables (RSFC/cognitive ability, RSFC/psychopathology, cortical thickness/cognitive ability, cortical thickness/psychopathology). For each imaging modality and each behavioral phenotype, we selected the 99th percentile (largest 1%) of brain-wide associations in the full discovery set (Extended Data Fig. S9). To explore how these estimates varied as a function of sample size, models were trained on subsamples of the discovery dataset at logarithmically spaced sample sizes (RSFC: N=25,33,45, 60, 80, 100, 145, 200, 256, 350, 460, 615, 825, 1,100, 1,475, 1,964; cortical thickness: N=25,33,45, 60, 80, 100, 145, 200, 256, 350, 460, 615, 825, 1,100, 1,475, 1,814) and tested on the full replication set (N=1,790). In order to model sampling variability as a function of sample size, 100 bootstrap (sampling with replacement) samples were generated for each sample size. To generate an out-of-sample replication coefficient (R^2^), we first determined the out-of-sample (replication set) sum of squares from the behavioral data (mean squared subtracted from sum). Next, a regression model linking the subsample of the discovery set brain and behavioral data (e.g., cognitive ability) was fit. The resulting best fit regression model (slope and intercept), were applied to replication brain data and subtracted from the out-of-sample behavior to obtain the residuals. We then determined the sum-of-squares of the residuals by subtracting the mean squared from the sum, dividing these sum-of-squared residuals by the original sum-of-squares, and subtracting them from 1 to obtain the out-of-sample replication coefficient (R^2^).

### Multivariate Replication: Support Vector Regression (SVR)

Support vector regression (SVR) with a linear kernel was performed using the e1071 package in the R environment (version 3.5.2) to predict primary behaviors (psychopathology, cognitive ability) and other demographics and behavioral phenotypes (Extended Data Fig. S14) from individual differences in either RSFC or cortical thickness. One hundred bootstrap samples (sampling with replacement) were generated for each sample size. Hyperparameter tuning was examined in 1) split halves of the full discovery sample for multiple cognitive (NIH Toolbox) and psychopathology (CBCL) behaviors and 2) 10-fold cross-validation within the full discovery sample for primary behaviors (psychopathology, cognitive ability; Extended Data Fig. S10 and S11), but did not appreciably change out-of-sample prediction estimates to the replication sample (e.g., average out-of-sample correlation difference between tuned and non-tuned models: RSFC = −0.006, Cortical Thickness = 0.014; Extended Data Fig. S10,S11). Figure 3a,c use default hyperparameters and PCA dimensionality reduction (with a threshold of 50% variance explained in the discovery set, for each sample size) prior to SVR, given that this procedure balanced out-of-sample prediction and model complexity for nearly all model types (Extended Data Fig. S10,11). Replication set data were not used to estimate principal components, but rather replication set data were projected into component space via independently estimated loading matrices for each subsample of the discovery set to prevent bias. An alternative strategy of univariate feature ranking was also examined, where SVR models were trained on the 5,000, 10,000, or 15,000 vertices (cortical thickness) or edges (RSFC) with the highest correlation to the variable of interest in the training dataset, but this approach resulted in lower out-of-sample prediction (Extended Data Fig. S10 and S11). Significance thresholds for out-of-sample replication were estimated via permutation testing (1,000 iterations) with models trained on the full discovery set (RSFC: N=1964; cortical thickness: N=1,814) and tested on the full replication set.

### Multivariate Replication: Canonical Correlation Analysis (CCA)

Canonical correlation analysis (CCA) was performed using Matlab’s (2019A) *cannoncor.m* function to predict the NIH Toolbox and CBCL from individual differences in either RSFC or cortical thickness. Equivalent bootstrapping and subsampling of the discovery set were tested and applied to the replication set, as in the SVR analyses. In order to model sampling variability across sample sizes, 100 bootstrap (sampling with replacement) samples were generated for each sample size. As with SVR, Figure 3b,d used principal-component analysis (PCA) dimensionality reduction (threshold of 20% variance explained in the discovery set, for each sample size) prior to CCA given that this maximized out-of-sample prediction (Extended Data Fig. S12,S13). CCA models were fit on iteratively larger subsamples of the discovery (in-sample) data set. The first canonical vector was extracted and applied to the full replication (out-of-sample) brain data to predict replication set (out-of-sample) behavior. Prediction accuracy was quantified using Pearson r, expressing the correlation between the matrix products of the first canonical vector (from the discovery set) and replication brain and behavioral data. Significance thresholds for out-of-sample replication were estimated via permutation testing (1,000 iterations) with models trained on the full discovery set (RSFC: N=1964; cortical thickness: N=1,814) and tested on the full replication set.

### Towards A New Era of Brain-Wide Association Studies (BWAS)

In Fig. 4, sampling variability, statistical errors (false positives, false negatives, inflation, sign errors), and out-of-sample replication were plotted on a common scale as a function of sample size (y-axis: 0-1 for sampling variability [r], 0-100% for statistical errors [cumulative sum across all four error types], 0-100% for out-of-sample replication [r]). Out-of-sample replication (r) was normalized by the maximum in-sample (discovery) correlation by dividing the mean replicated coefficient (r) across subsamples by the maximum in-sample (discovery) correlation. All three curves (sampling variability, statistical errors, and out-of-sample replication) were based on the largest univariate and multivariate brain-wide association (RSFC and cognitive ability).

## Acknowledgements

We thank Anders M. Dale, Terry L. Jernigan, Weiqi Zhao, Carolina Makowski, Chun Chieh Fan, and Clare Palmer for their thoughtful comments on the manuscript. This work was supported by NIH grants MH100019 (S.M.), MH121518 (S.M.), DA007261 (D.F.M.), NS090978 (B.P.K.), DA007261 (A.S.H.), NS110332 (D.J.N.), NS115672 (A.Z.), MH112473 (T.O.L.), MH104592 (D.J.G.), DA041148 (D.A.F.), DA04112 (D.A.F.), MH115357 (D.A.F.), MH096773 (D.A.F, N.U.F.D.), MH122066 (D.A.F., N.U.F.D.), MH121276 (D.A.F., N.U.F.D), MH124567 (D.A.F., N.U.F.D.), NS088590 (N.U.F.D.), and the Andrew Mellon Predoctoral Fellowship (B.T.-C.), Lynne and Andrew Redleaf Foundation (D.A.F.), Kiwanis Neuroscience Research Foundation (N.U.F.D.), the Jacobs Foundation grant 2016121703 (N.U.F.D.).

## ABCD acknowledgement

Data used in the preparation of this article were obtained from the Adolescent Brain Cognitive Development (ABCD) Study (https://abcdstudy.org), held in the NIMH Data Archive (NDA). This is a multisite, longitudinal study designed to recruit more than 10,000 children age 9-10 and follow them over 10 years into early adulthood. The ABCD Study is supported by the National Institutes of Health and additional federal partners under award numbers U01DA041022, U01DA041028, U01DA041048, U01DA041089, U01DA041106, U01DA041117, U01DA041120, U01DA041134, U01DA041148, U01DA041156, U01DA041174, U24DA041123, U24DA041147, U01DA041093, and U01DA041025. A full list of supporters is available at https://abcdstudy.org/federal-partners.html. A listing of participating sites and a complete listing of the study investigators can be found at https://abcdstudy.org/scientists/workgroups/. ABCD consortium investigators designed and implemented the study and/or provided data but did not necessarily participate in analysis or writing of this report. This manuscript reflects the views of the authors and may not reflect the opinions or views of the NIH or ABCD consortium investigators.

## XSEDE and Pittsburgh Supercomputing Center Acknowledgement

This work used the Extreme Science and Engineering Discovery Environment (XSEDE), which is supported by National Science Foundation grant number ACI-1548562. Specifically, it used the Bridges system, which is supported by NSF award number ACI-1445606, at the Pittsburgh Supercomputing Center (PSC^6465^*TG-IBN200009*).

## Authorship Contributions

### Conception

S.M., B.T.-C., D.A.F., N.U.F.D.;

### Design of the work

S.M., B.T.-C., F.J.C., D.F.M., B.T.T.Y., B.L., D.A.F., N.U.F.D.;

### Acquisition, analysis, interpretation of data

S.M., B.T.-C., F.J.C., D.F.M., B.P.K., A.S.H., M.R.D., W.F., R.L.M., E.F., O.M.-D, A.M.G., E.A.E., A.J.P., M.C., O.D., L.A.M., G.C., J.U., K.S., T.O.L., W.K.T., D.J.G., S.E.P., T.E.N., B.T.T.Y., D.M.B., H.G., B.L., D.A.F., N.U.F.D.;

### Manuscript writing and revising

S.M., B.T.-C., F.J.C., D.F.M., B.P.K., A.S.H., M.R.D., W.F., R.L.M., E.F., O.M.-D., A.M.G., E.A.E., A.J.P., M.C., O.D., L.A.M., G.C., J.U., K.S., A.T., J.C., D.J.N., A.Z., N.A.S., A.N.V., T.O.L., W.K.T., D.J.G., S.E.P., T.E.N., B.T.T.Y., D.M.B., H.G., B.L., D.A.F., N.U.F.D.

## Code Sharing

Analysis code specific to this study can be found here: https://gitlab.com/DosenbachGreene/.

## Data Availability

The ABCD data repository grows and changes over time. The ABCD data used in this report came from ABCD collection 3165 and the Annual Release 2.0, DOI 10.15154/1503209.

---

**Table S1.**
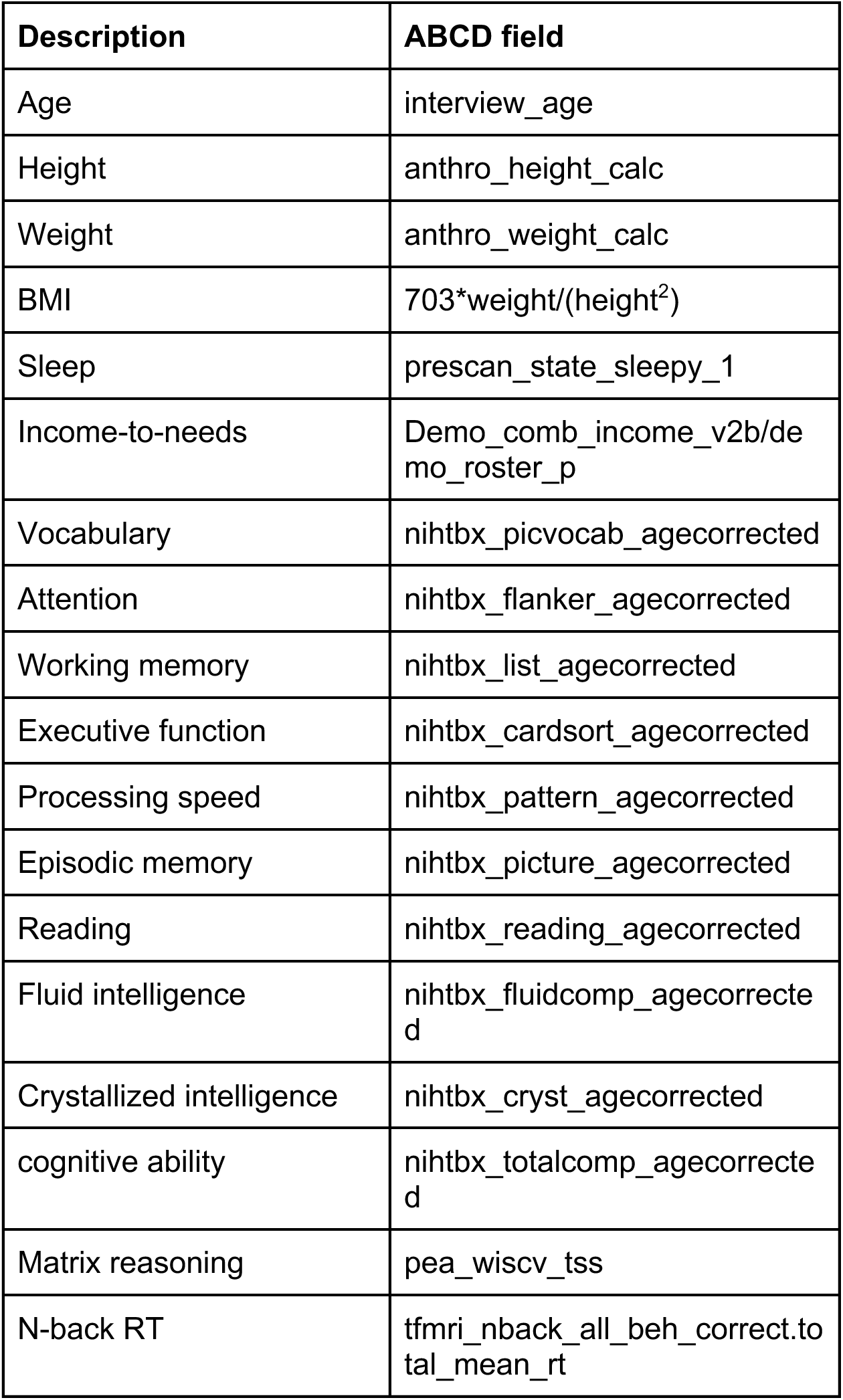
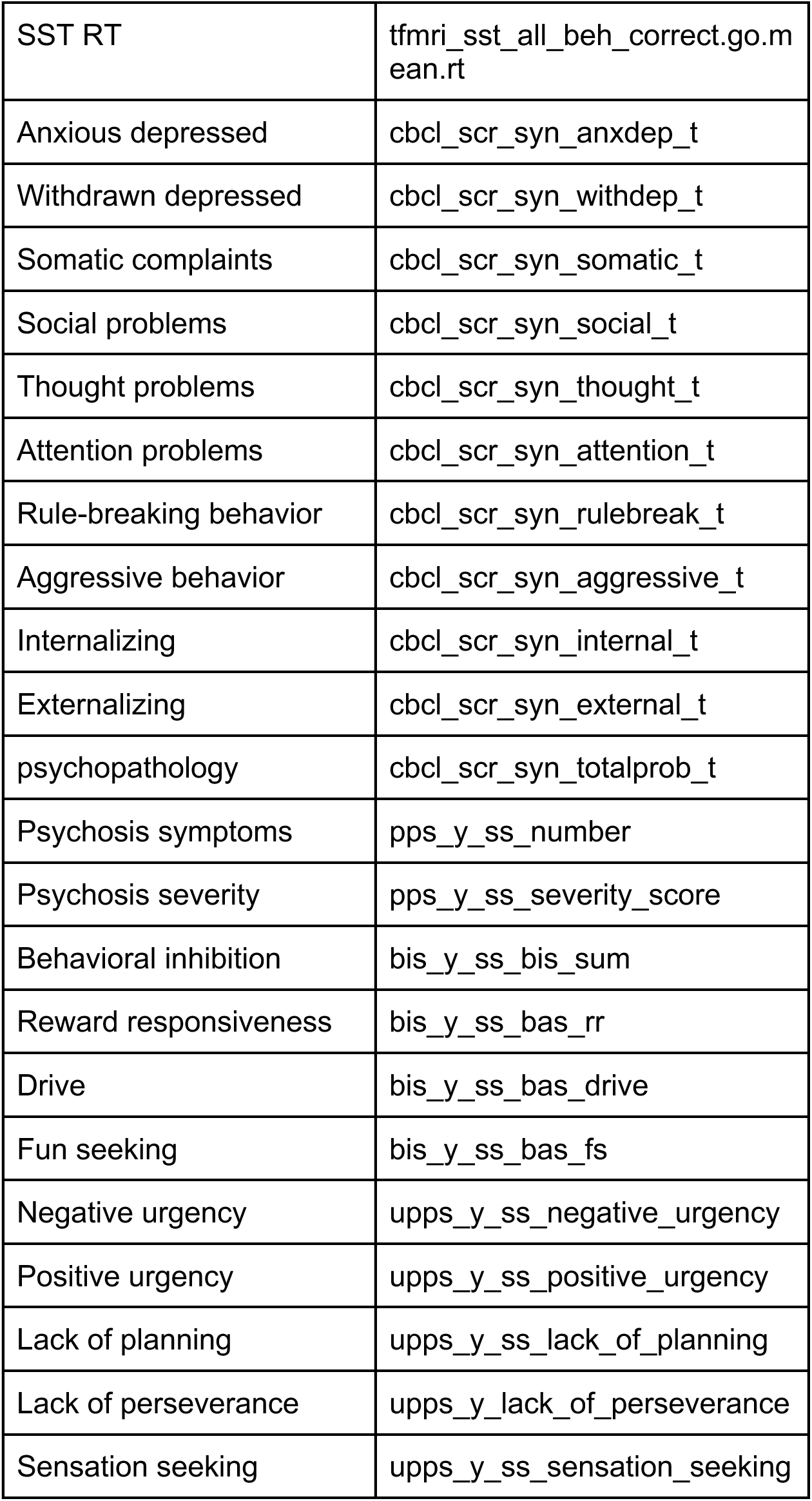
Lookup table showing the original ABCD variable names with the corresponding descriptive labels used in the manuscript. More details on the demographic and behavioral measures can be found in the ABCD data dictionary.

**Extended Data Fig S1.**
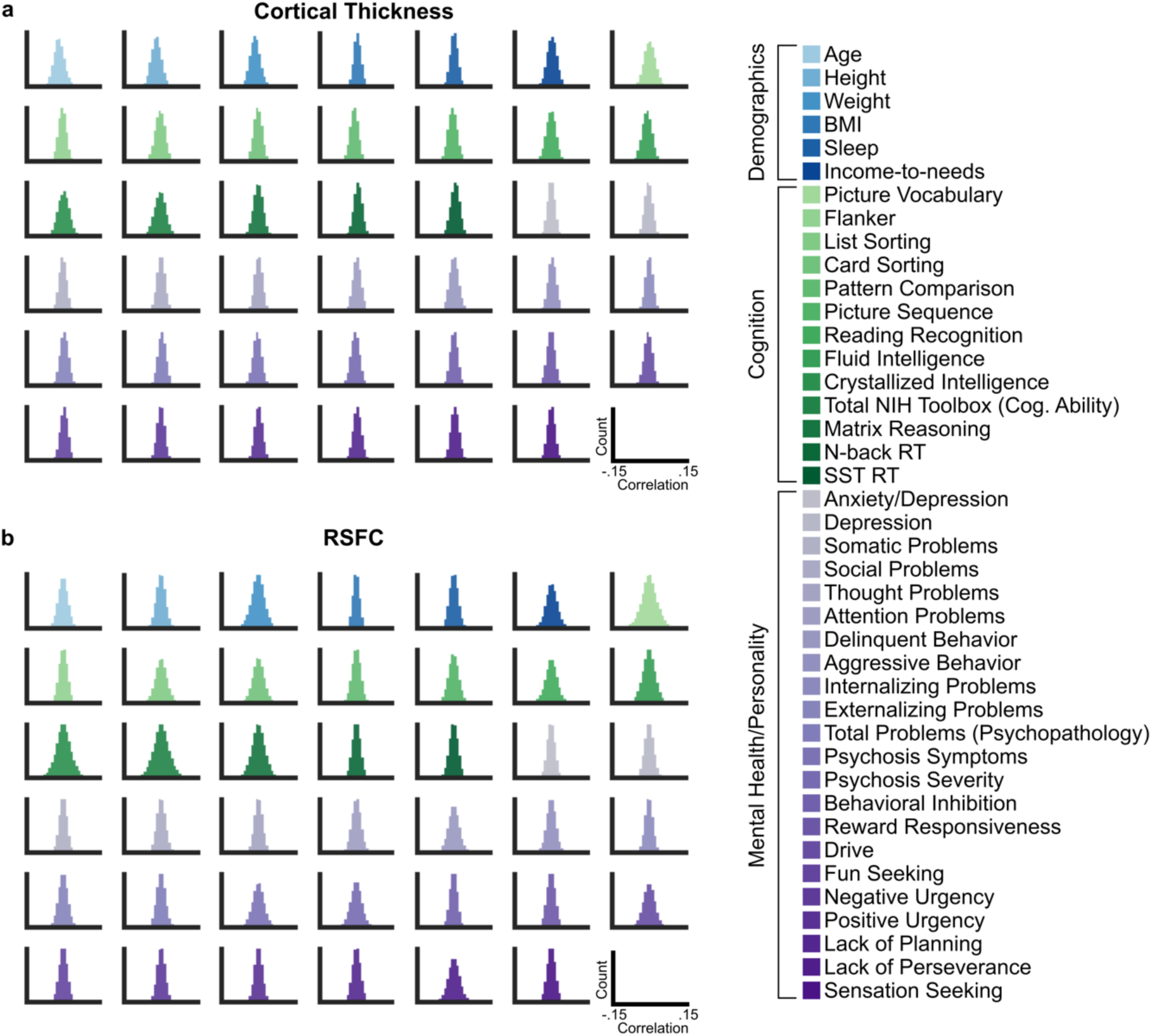
Histograms of all **(a)** cortical thickness association with demographics and behavioral phenotypes and **(b)** RSFC associations with demographics and behavioral phenotypes. For each demographic and behavioral (cognition, mental health) measure (N=41). Correlations with brain measures were generated across multiple levels of scale (cortical thickness: vertices, ROIs, networks; RSFC: ROI-ROI pairs (edges), principal components, networks). The ordering of subgraphs follows the ordering of measures in the legend (e.g., the most top-left panel shows sampling variability between Age and every other demographic and behavioral measure).

**Extended Data Fig. S2.**
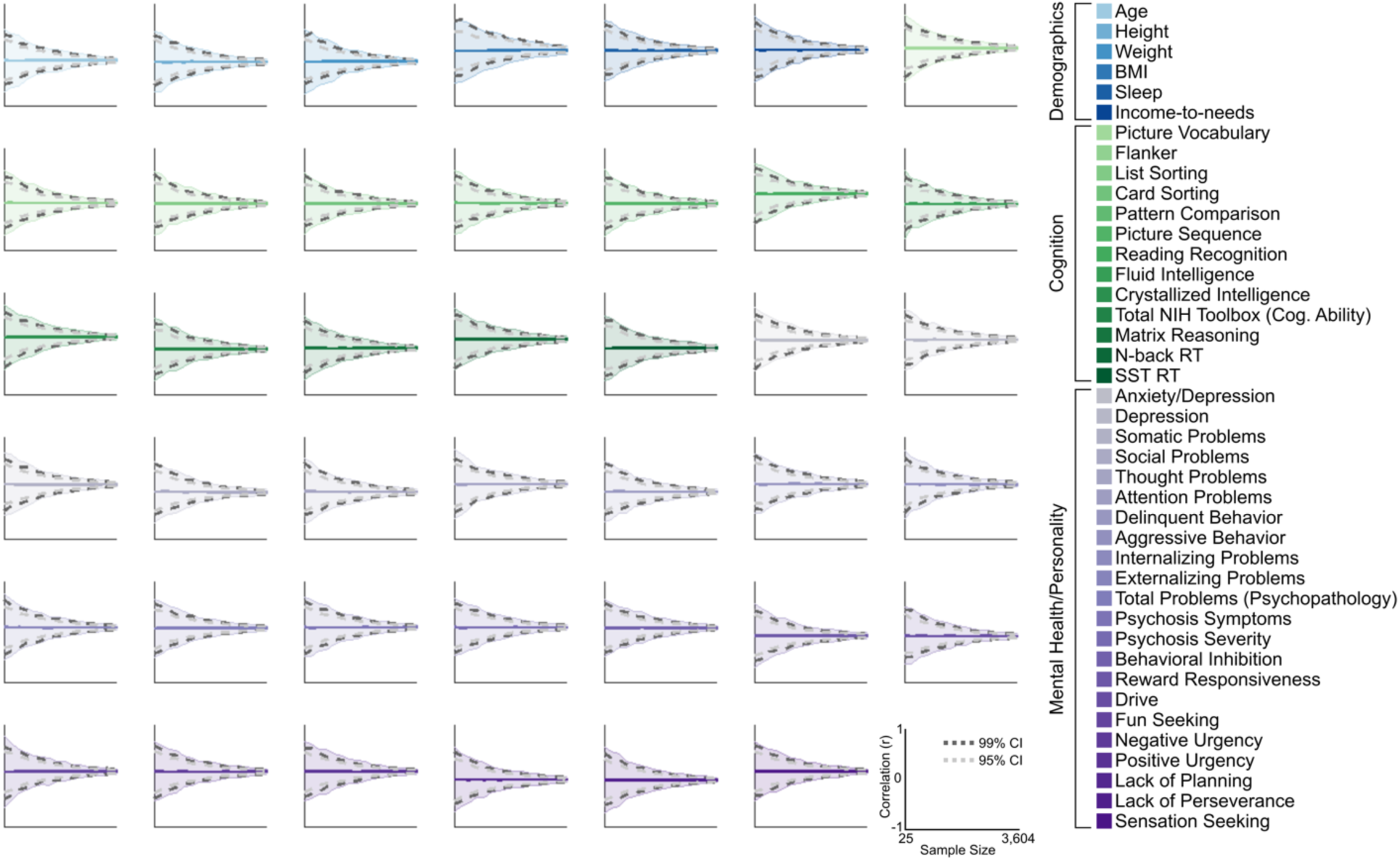
Sampling variability of the correlation between cortical thickness and each demographic, cognitive and mental health measure (N=41). For each brain-wide association, 16,000 resampled studies (1,000 subsamples for each sample size) were generated. Across sample sizes, sampling variability of the largest brain-wide association is depicted as the range of observable correlations (shaded area), as well as the 99% confidence interval (dark gray) and 95% confidence interval (light gray). For each sampling bin, the dark colored line represents the mean brain-wide association across the 1,000 resamples. The ordering of subgraphs follows the ordering of measures in the legend (e.g., the most top-left panel shows sampling variability between Age and every other demographic and behavioral measure).

**Extended Data Fig. S3.**
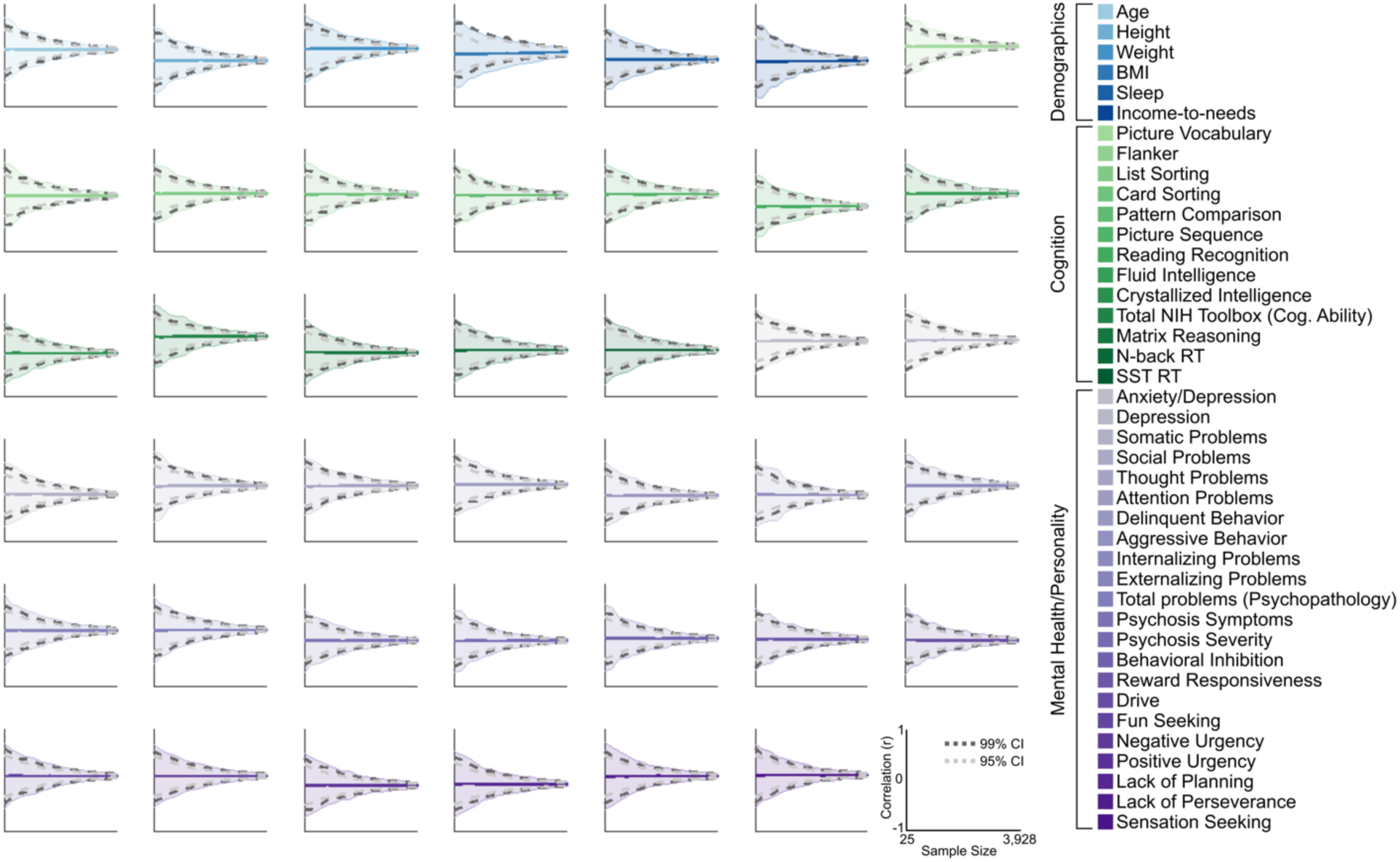
Sampling variability of the correlation between RSFC and each demographic, cognitive and mental health measure (N=41). For each brain-wide association, 16,000 resampled studies (1,000 subsamples for each sample size) were generated. Across sample sizes, sampling variability of the largest brain-wide association is depicted as the range of observable correlations (shaded area), as well as the 99% confidence interval (dark gray) and 95% confidence interval (light gray). For each sampling bin, the dark colored line represents the mean brain-wide association across the 1,000 resamples. The ordering of subgraphs follows the ordering of measures in the legend (e.g., the most top-left panel shows sampling variability between Age and every other demographic and behavioral measure).

**Extended Data Fig. S4.**
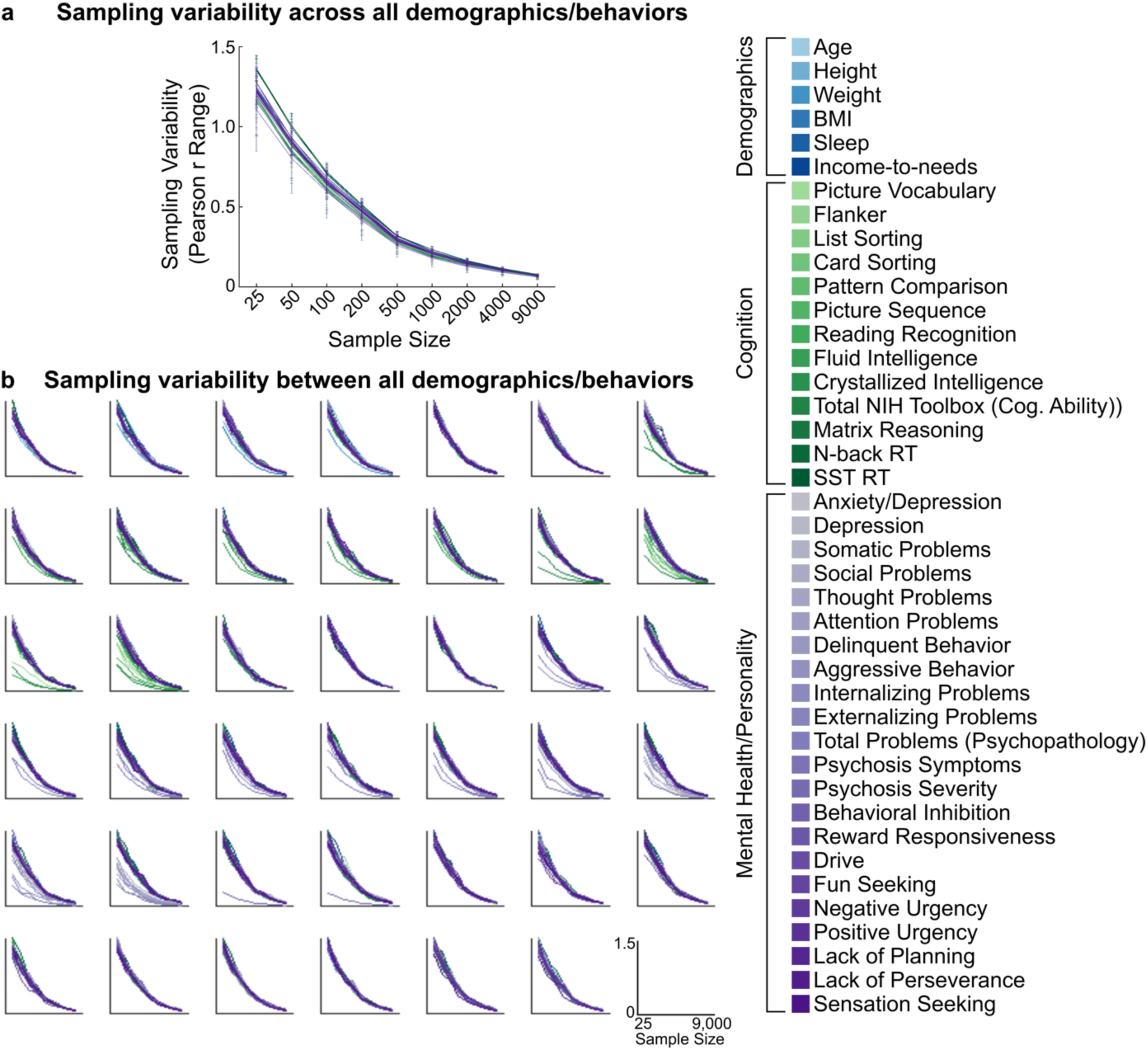
Sampling variability of associations between demographic and behavioral measures. **(a)** Sampling variability (range in observable correlations) as a function of sample size. Each line represents the average sampling variability for the correlation between a measure (demographic, cognition, mental health) and all other measures. Error bars denote one standard deviation across all measures. Note that sampling variability ranges from 0-2 for Pearson correlations, given observables correlations range from −1 to 1. **(b)** Each subgraph depicts sampling variability as a function of sample size for a given demographic or behavioral measure (N=41) to every other demographic/behavioral measure. The ordering of subgraphs follows the ordering of measures in the legend (e.g., the most top-left panel shows sampling variability between Age and every other demographic and behavioral measure). Each curve is colored corresponding to the measures in the figure legend.

**Extended Data Fig. S5.**
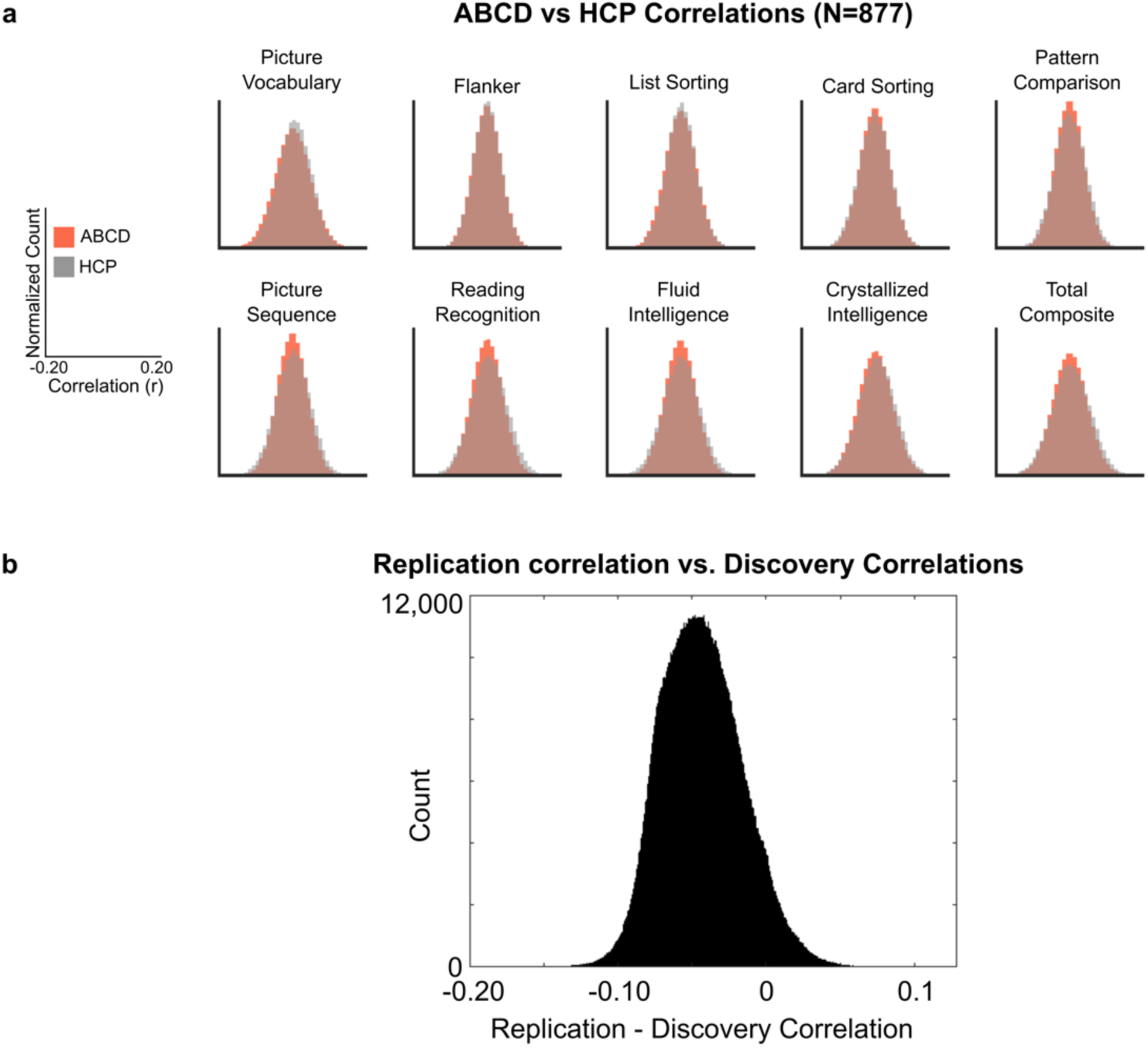
**(a)** Effect sizes observed in subsamples of ABCD (N=877) replicate full sample brain-phenotype correlations from HCP (N=877). ABCD data were subsampled to match the sample size of HCP (N=877). Data were subsampled 100 times. For each NIH Toolbox subscale the correlation between every ROI pair and behavior was generated for HCP data and each of the 100 resamples of ABCD data. Each histogram (ABCD: red; HCP: gray) was generated from these ROI-ROI pair brain-behavioral phenotype associations. **(b)** Difference in replication correlations (replication set - discovery set). The top 1% largest effect sizes were determined for each demographic/behavioral phenotype across 100 resampled discovery data sets (N=1,964 in each) and compared (subtracted) to a replication set (N-1,964). On average, replicated correlations were smaller (r=0.043) than the in-sample discovery set, indicating that, on average, correlations with ~2,000 individuals are inflated.

**Extended Data Fig S6.**
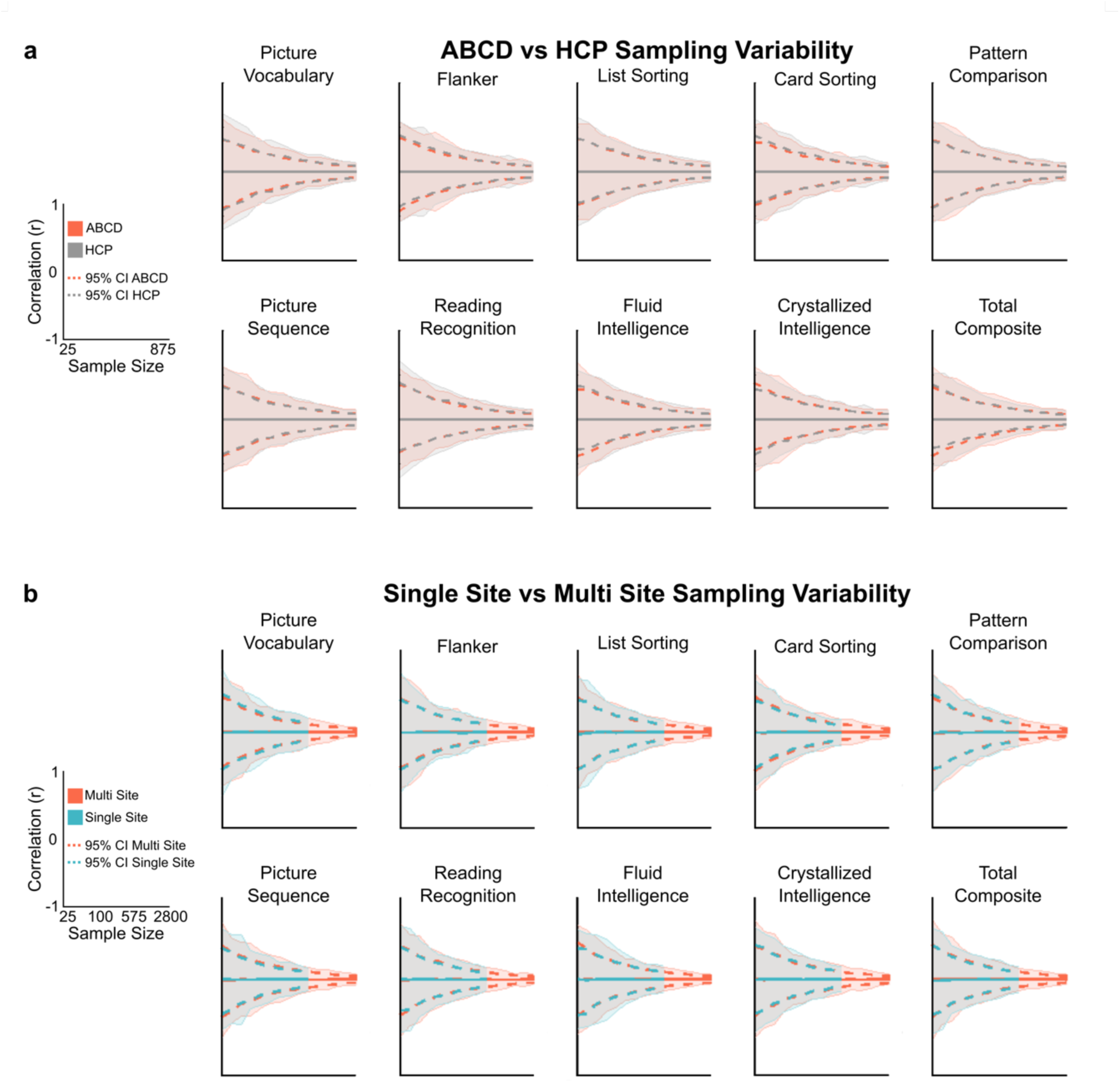
**(a)** Sampling variability RSFC associations with the NIH Toolbox in equally sized samples (N=877) from HCP (gray) and ABCD (red). **(b)** Sampling variability of RSFC association with the NIH Toolbox in a single-site ABCD sample (site 16; N=603; blue) and every other ABCD site (N=3,325; red).

**Extended Data Fig S7.**
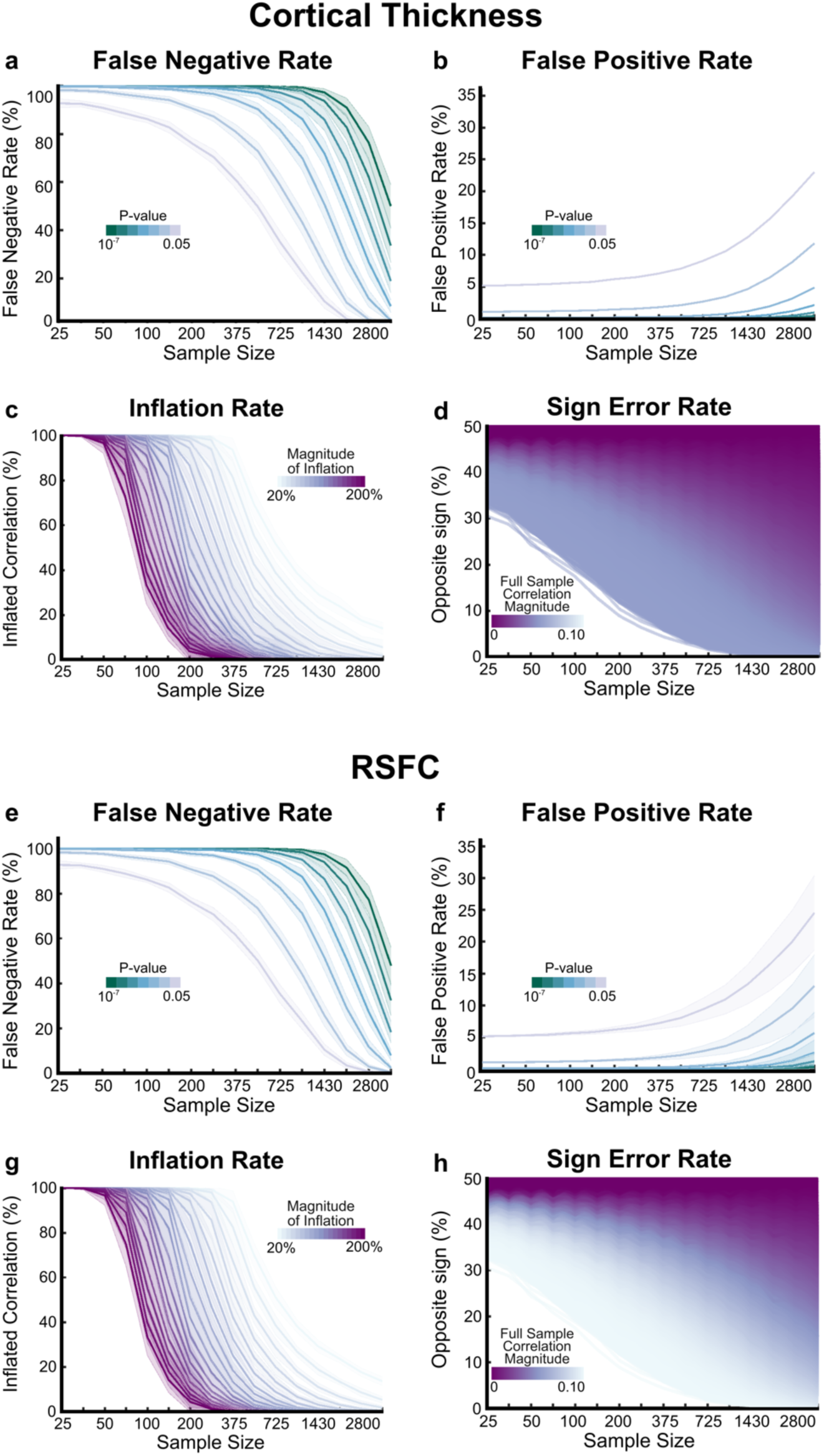
Statistical errors for univariate associations between cortical thickness and all 41 demographic and behavioral measures. **(a)** False negative rate in RSFC (edgewise) correlations with demographics and behavioral phenotypes varying as a function of p-value. The most stringent p-value (10^−7^) is equivalent to a p<0.05 Bonferroni corrected across all ROI-ROI pairs, whereas p<0.05 is equivalent to no multiple comparisons correction. **(b)** False positive rate in RSFC correlations with behavioral phenotypes varying as a function of p-value. **(c)** Inflation rate of the cortical thickness correlation with behavioral phenotypes, depicting the probability of observing an inflated correlation as a function of sample size. Color gradient represents the magnitude of inflation. **(d)** Probability of observing the opposite sign of the correlation observed between cortical thickness and behavioral phenotypes in the full sample (N=3,928) as a function of sample size. Color gradient represents the effect size of the correlation in the full sample. For each panel, darker lines represent the mean across the 41 behavioral phenotypes and shaded error bars represent one standard deviation from the mean. **(e-h)** RSFC associations with demographics and behavioral phenotypes for each statistical error.

**Extended Data Fig S8.**
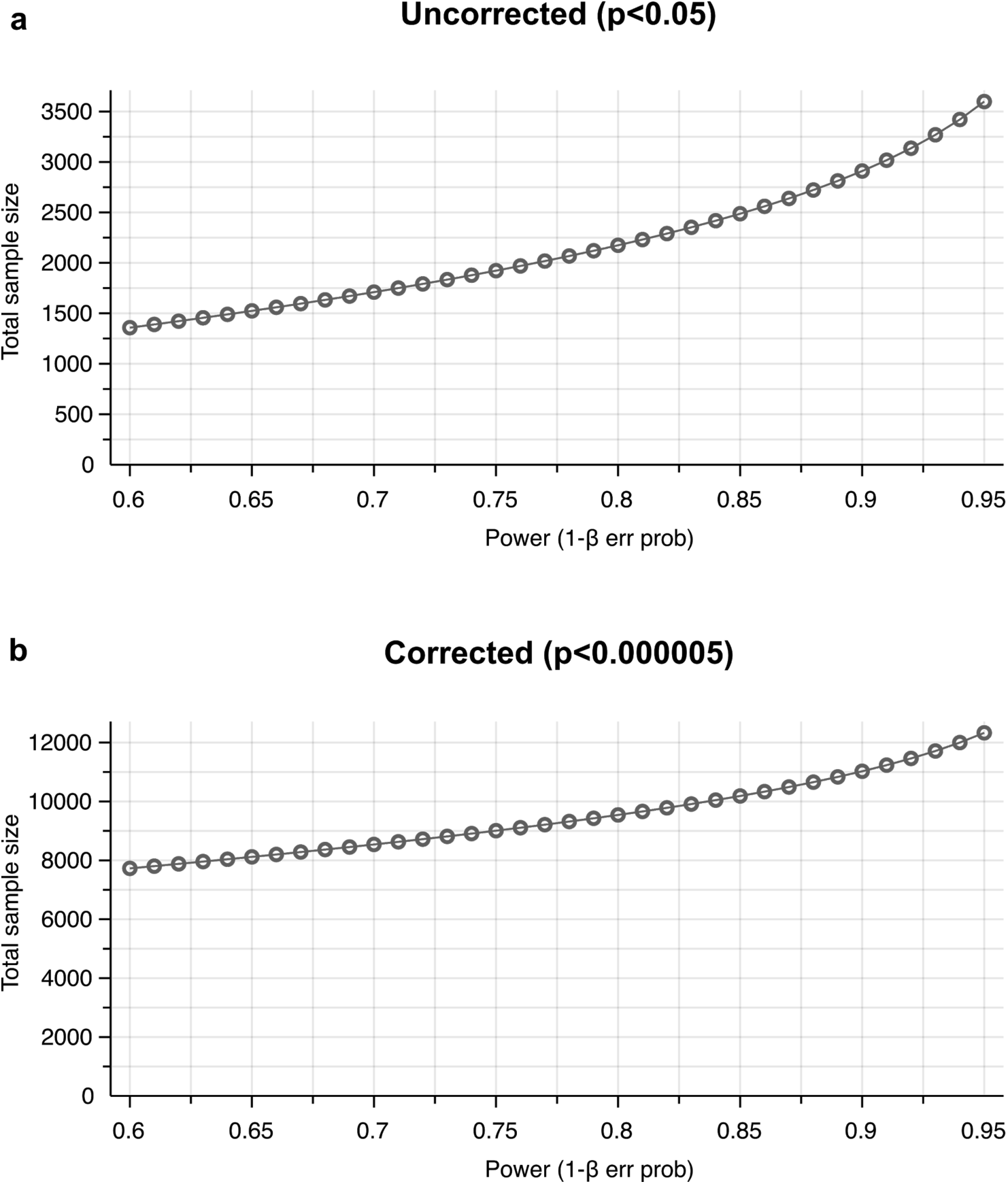
Analytic requisite sample sizes to detect 99th percentile (largest 1%; r=0.06) univariate brain-wide associations at **(a)** p<0.05 (uncorrected) and **(b)** p<10^−7^ (p<0.05 Bonferroni corrected).

**Extended Data Fig. S9.**
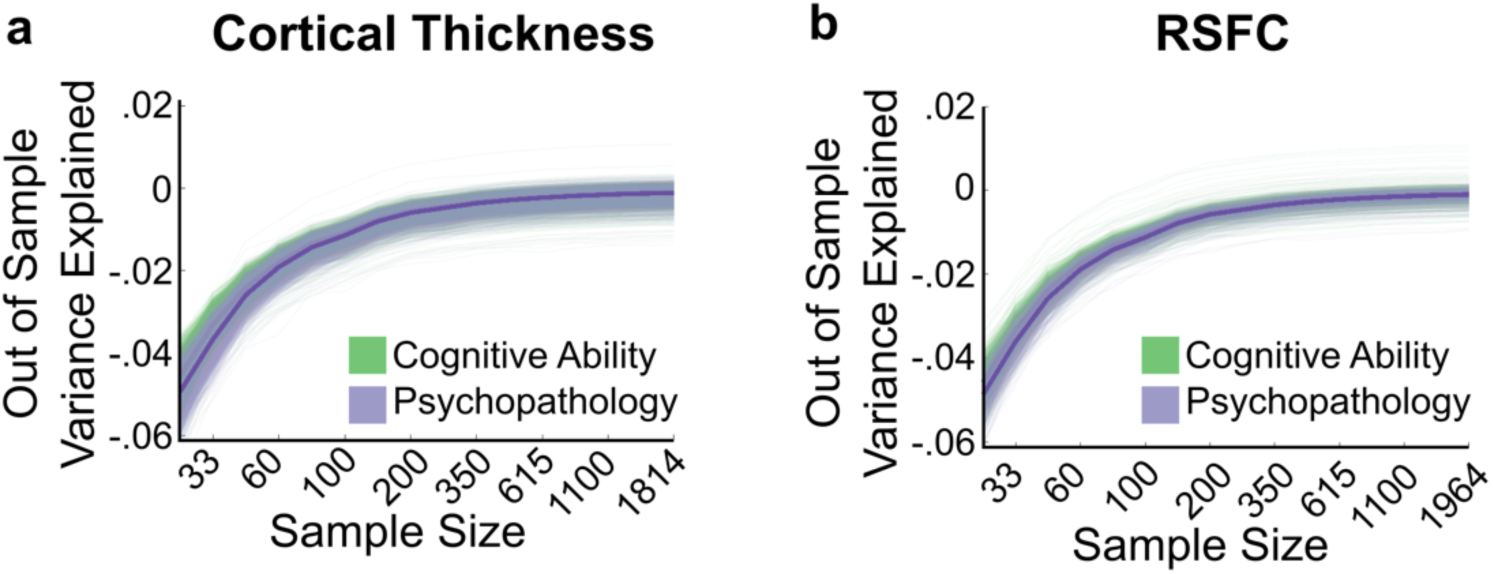
Univariate out-of-sample replication (R^2^) for **(a)** cortical thickness and **(b)** RSFC relationships with cognitive ability and psychopathology across sample sizes.

**Extended Data Fig. S10.**
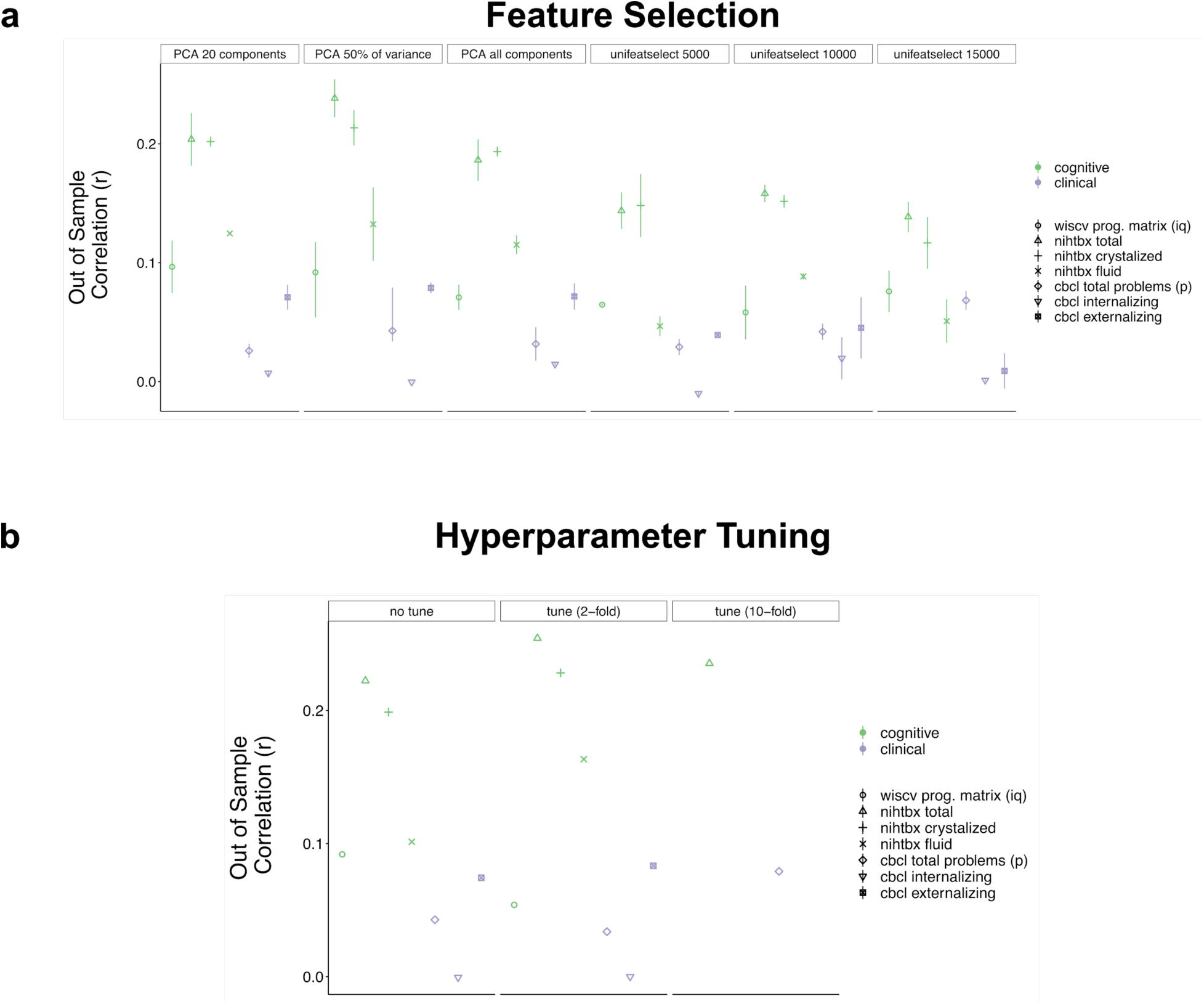
Out-of-sample correlation (r) for cortical thickness associations with behavior trained on the full discovery set (N=1,964) and tested on the full replication set (N=1,790) using support vector regression (SVR) across **(a)** feature selection procedure (principal components left, univariate feature ranking right) and **(b)** hyperparameter tuning. Note, point range in (a) shows variability across tuned and non-tuned models from (b); (b) displays tuned and non-tuned models using PCA with 50% of the variance retained.

**Extended Data Fig. S11.**
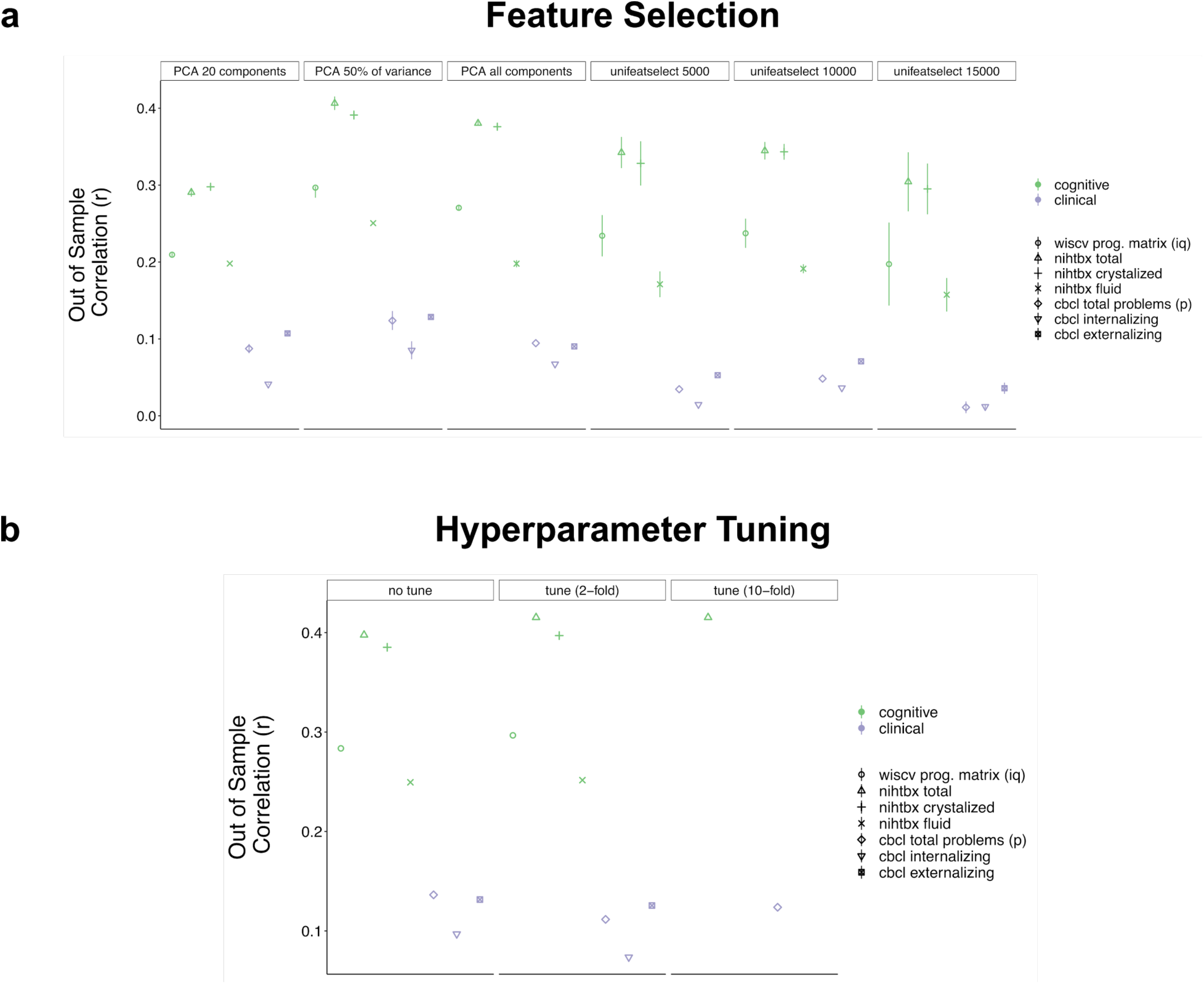
Out-of-sample correlation (r) for RSFC associations with behavior trained on the full discovery set (N=1,964) and tested on the full replication set (N=1,790) using support vector regression across **(a)** feature selection procedure (principal components left, univariate feature ranking right) and **(b)** hyperparameter tuning. Note, point range in (a) shows variability across tuned and non-tuned models from (b); (b) displays tuned and non-tuned models using PCA with 50% of the variance retained.

**Extended Data Fig. S12.**
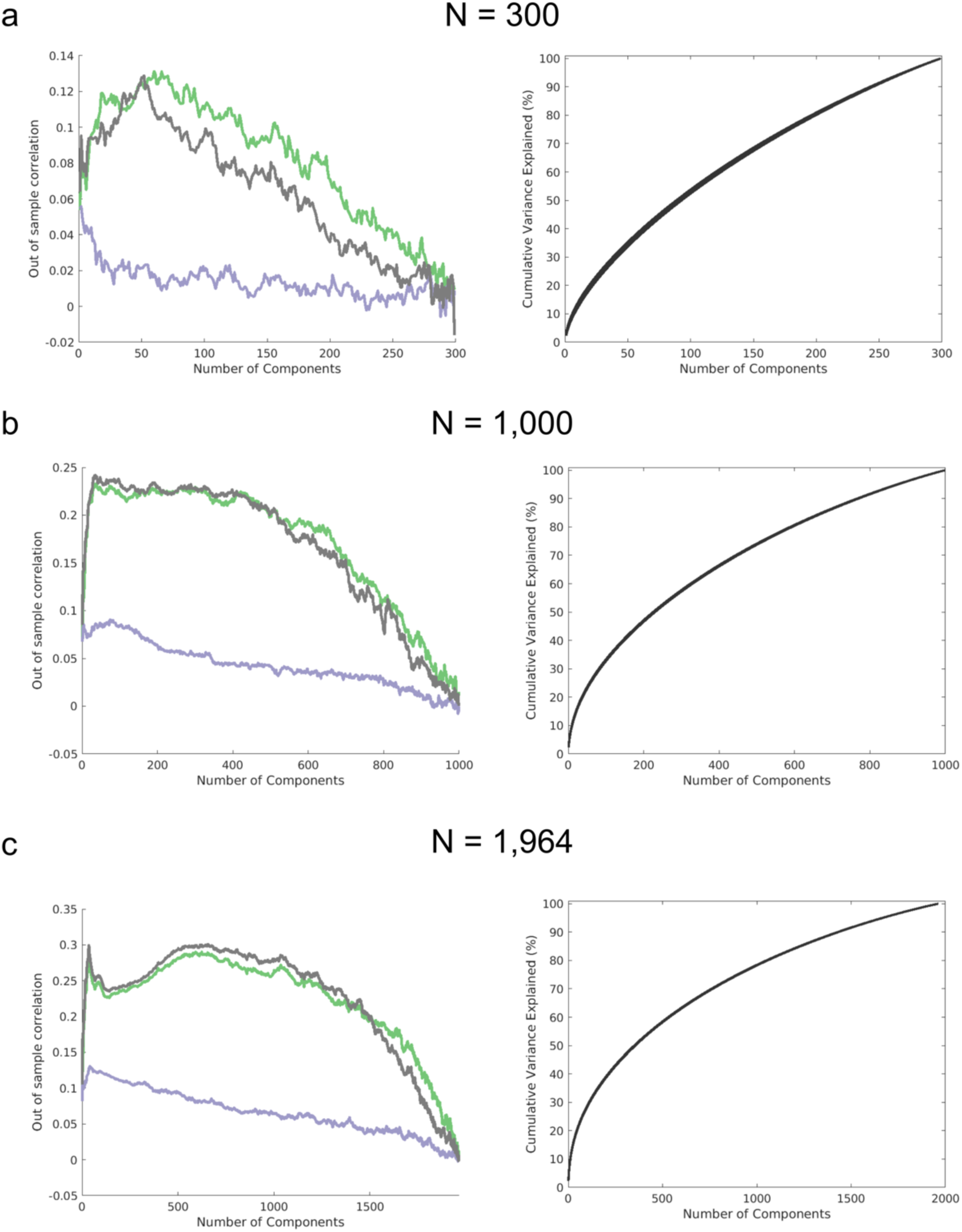
Out-of-sample replication (r) as a function of the number of principal components included in the RSFC CCA models. Across three distinct sample sizes (N=300, N=1,000, and N=1,964; 100 iterations of each), ~20% of the cumulative principal component variance maximized the out-of-sample correlation.

**Extended Data Fig. S13.**
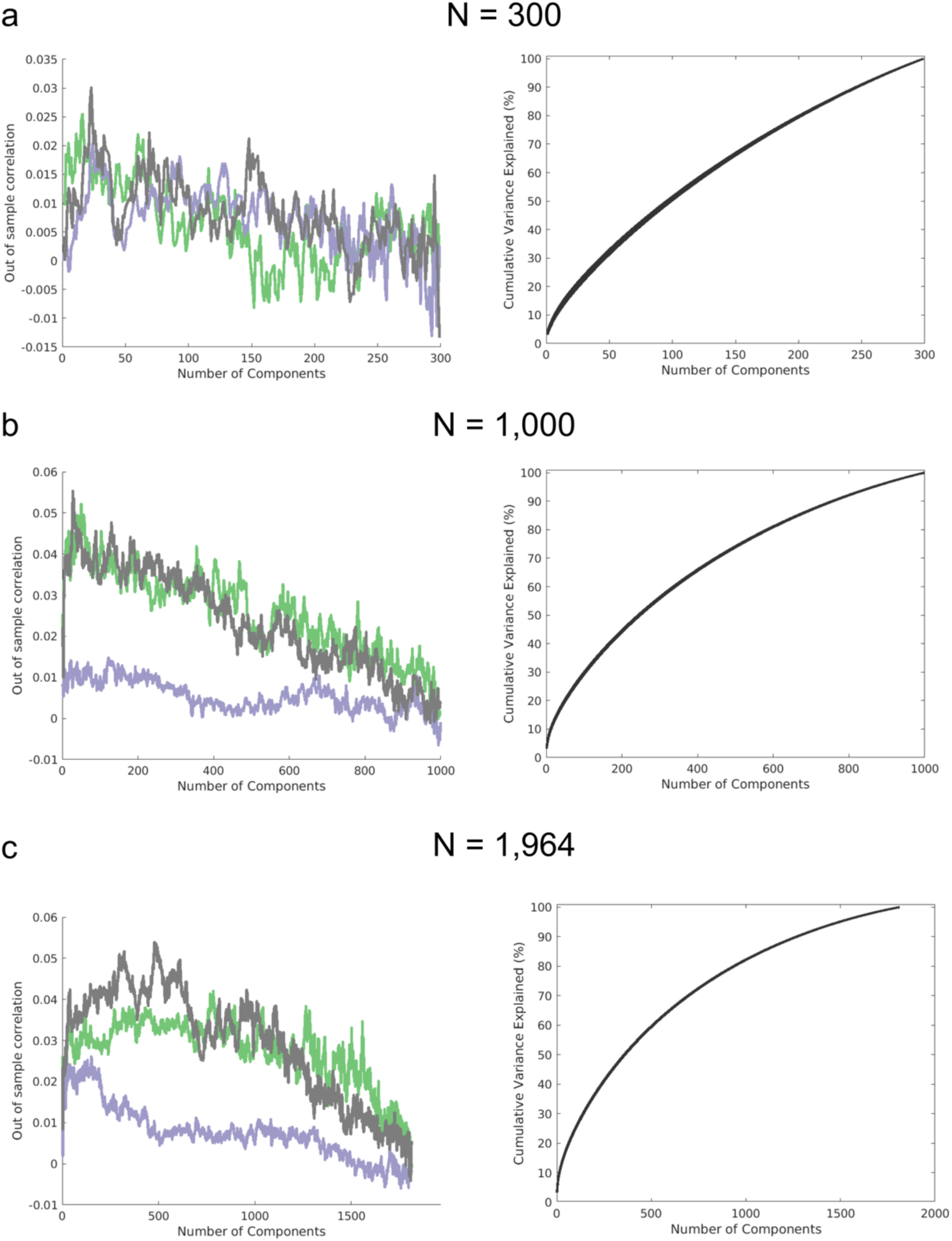
Out-of-sample replication (r) as a function of the number of principal components included in the cortical thickness CCA models. Similar to RSFC, across three distinct sample sizes (N=300, N=1,000, and N=1,964; 100 iterations of each), ~20% of the cumulative principal component variance maximized the out-of-sample correlation.

**Extended Data Fig. S14.**
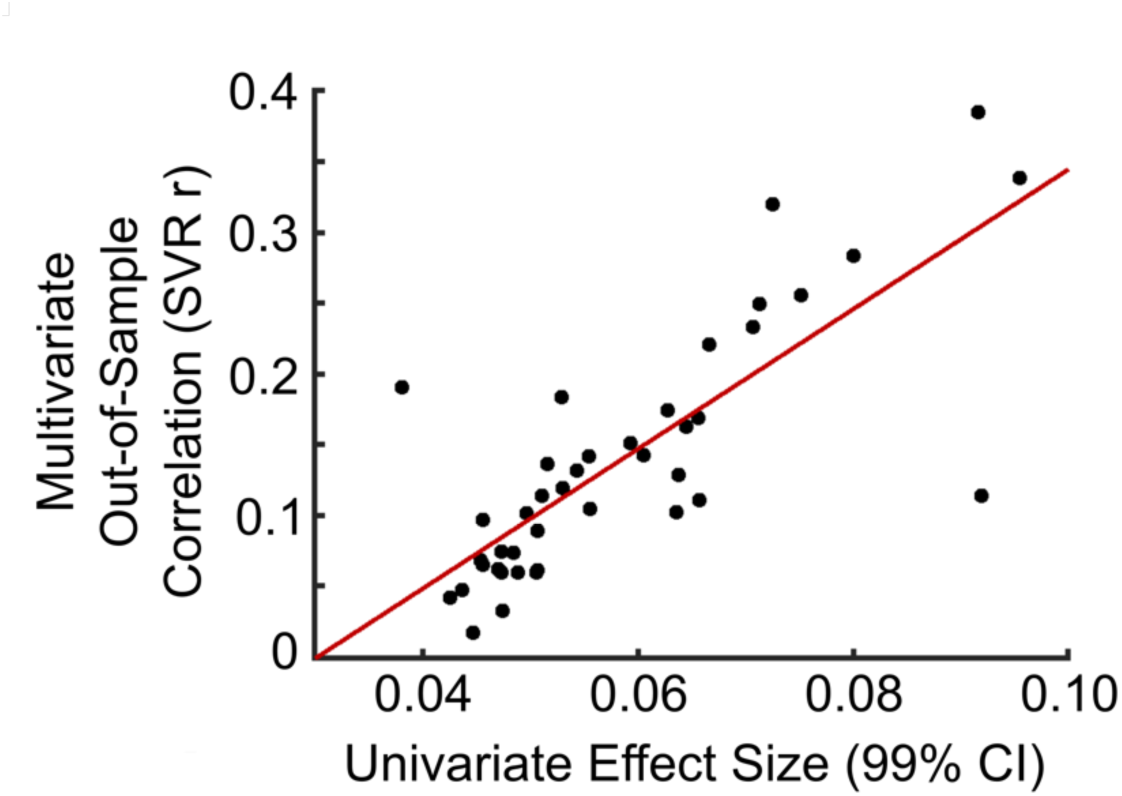
Multivariate (SVR) out-of-sample replication (r) as a function of the univariate RSFC-phenotype correlations (99% confidence interval) across the 41 behavioral phenotypes. Multivariate out-of-sample replication was positively correlated with univariate effect sizes (r=0.79, p<0.001).

## References

1. Poldrack, R. A. et al. Scanning the horizon: towards transparent and reproducible neuroimaging research. Nat. Rev. Neurosci. 18, 115–126 (2017).

2. Szucs, D. & Ioannidis, J. P. Sample size evolution in neuroimaging research: an evaluation of highly-cited studies (1990-2012) and of latest practices (2017-2018) in high-impact journals. Neuroimage 117164 (2020).

3. Volkow, N. D. et al. The conception of the ABCD study: From substance use to a broad NIH collaboration. Dev. Cogn. Neurosci. 32, 4–7 (2018).

4. Button, K. S. et al. Power failure: why small sample size undermines the reliability of neuroscience. Nat. Rev. Neurosci. 14, 365–376 (2013).

5. Ioannidis, J. P. A., Munafò, M. R., Fusar-Poli, P., Nosek, B. A. & David, S. P. Publication and other reporting biases in cognitive sciences: detection, prevalence, and prevention. Trends Cogn. Sci. 18, 235–241 (2014).

6. Botvinik-Nezer, R. et al. Variability in the analysis of a single neuroimaging dataset by many teams. Nature 582, 84–88 (2020).

7. Dale, A. M., Fischl, B. & Sereno, M. I. Cortical Surface-Based Analysis. NeuroImage vol. 9 179–194 (1999).

8. Sowell, E. R., Thompson, P. M., Holmes, C. J., Jernigan, T. L. & Toga, A. W. In vivo evidence for post-adolescent brain maturation in frontal and striatal regions. Nat. Neurosci. 2, 859–861 (1999).

9. Raichle, M. E. et al. A default mode of brain function. Proc. Natl. Acad. Sci. U. S. A. 98, 676–682 (2001).

10. Dosenbach, N. U. F. et al. A core system for the implementation of task sets. Neuron 50, 799–812 (2006).

11. Masouleh, S. K., Eickhoff, S. B., Hoffstaedter, F., Genon, S. & Alzheimer’s Disease Neuroimaging Initiative. Empirical examination of the replicability of associations between brain structure and psychological variables. eLife vol. 8 (2019).

12. Drysdale, A. T. et al. Resting-state connectivity biomarkers define neurophysiological subtypes of depression. Nat. Med. 23, 28–38 (2017).

13. Dinga, R. et al. Evaluating the evidence for biotypes of depression: Methodological replication and extension of. Neuroimage Clin 22, 101796 (2019).

14. Westlye, L. T., Grydeland, H., Walhovd, K. B. & Fjell, A. M. Associations between regional cortical thickness and attentional networks as measured by the attention network test. Cereb. Cortex 21, 345–356 (2011).

15. Boekel, W. et al. A purely confirmatory replication study of structural brain-behavior correlations. Cortex 66, 115–133 (2015).

16. Bishop, D. How scientists can stop fooling themselves over statistics. Nature vol. 584 9–9 (2020).

17. Nosek, B. A., Cohoon, J., Kidwell, M. & Spies, J. R. Estimating the Reproducibility of Psychological Science. doi:10.31219/osf.io/447b3.

18. Visscher, P. M. et al. 10 Years of GWAS Discovery: Biology, Function, and Translation. Am. J. Hum. Genet. 101, 5–22 (2017).

19. Begley, C. G. & Ellis, L. M. Drug development: Raise standards for preclinical cancer research. Nature 483, 531–533 (2012).

20. Schönbrodt, F. D. & Perugini, M. At what sample size do correlations stabilize? Journal of Research in Personality vol. 47 609–612 (2013).

21. Border, R. et al. No Support for Historical Candidate Gene or Candidate Gene-by-Interaction Hypotheses for Major Depression Across Multiple Large Samples. Am. J. Psychiatry 176, 376–387 (2019).

22. Farrell, M. S. et al. Evaluating historical candidate genes for schizophrenia. Mol. Psychiatry 20, 555–562 (2015).

23. Van Essen, D. C. et al. The WU-Minn Human Connectome Project: An overview. NeuroImage vol. 80 62–79 (2013).

24. Achenbach, T. M. Achenbach System of Empirically Based Assessment (ASEBA): Development, Findings, Theory, and Applications. (University of Vermont Research Center of Children Youth & Families, 2009).

25. Gershon, R. C. et al. NIH Toolbox for Assessment of Neurological and Behavioral Function. Neurology vol. 80 S2–S6 (2013).

26. Kanai, R. & Rees, G. The structural basis of inter-individual differences in human behaviour and cognition. Nat. Rev. Neurosci. 12, 231–242 (2011).

27. Szucs, D. & Ioannidis, J. P. A. Empirical assessment of published effect sizes and power in the recent cognitive neuroscience and psychology literature. PLoS Biol. 15, e2000797 (2017).

28. Gelman, A. & Carlin, J. Beyond Power Calculations: Assessing Type S (Sign) and Type M (Magnitude) Errors. Perspect. Psychol. Sci. 9, 641–651 (2014).

29. Marek, S. et al. Identifying reproducible individual differences in childhood functional brain networks: An ABCD study. Dev. Cogn. Neurosci. 40, 100706 (2019).

30. McIntosh, A. R. & Mišić, B. Multivariate Statistical Analyses for Neuroimaging Data. Annual Review of Psychology vol. 64 499–525 (2013).

31. Chen, J. et al. Shared and unique brain network features predict cognition, personality and mental health in childhood. doi:10.1101/2020.06.24.168724.

32. Sripada, C. et al. Prediction of neurocognition in youth from resting state fMRI. Mol. Psychiatry (2019) doi:10.1038/s41380-019-0481-6.

33. Smith, S. M. et al. A positive-negative mode of population covariation links brain connectivity, demographics and behavior. Nat. Neurosci. 18, 1565–1567 (2015).

34. Investigators, T. A. of U. R. P. & The All of Us Research Program Investigators. The ‘All of Us’ Research Program. New England Journal of Medicine vol. 381 668–676 (2019).

35. NCI-NHGRI Working> Group on Replication in Association Studies et al. Replicating genotype-phenotype associations. Nature 447, 655–660 (2007).

36. Lee, J. J. et al. Gene discovery and polygenic prediction from a genome-wide association study of educational attainment in 1.1 million individuals. Nat. Genet. 50, 1112–1121 (2018).

37. Lam, M. et al. RICOPILI: Rapid Imputation for COnsortias PIpeLIne. Bioinformatics 36, 930–933 (2020).

38. Palmer, C. E. et al. Determining the association between regionalisation of cortical morphology and cognition in 10,145 children. doi:10.1101/816025.

39. He, T. et al. Meta-matching: a simple framework to translate phenotypic predictive models from big to small data. doi:10.1101/2020.08.10.245373.

40. Zhao, W. et al. The Bayesian polyvertex score (PVS-B): a whole-brain phenotypic prediction framework for neuroimaging studies. doi:10.1101/813915.

41. Feczko, E. & Fair, D. A. Methods and Challenges for Assessing Heterogeneity. Biol. Psychiatry 88, 9–17 (2020).

42. Ciric, R. et al. Benchmarking of participant-level confound regression strategies for the control of motion artifact in studies of functional connectivity. NeuroImage vol. 154 174–187 (2017).

43. Dosenbach, N. U. F. et al. Real-time motion analytics during brain MRI improve data quality and reduce costs. Neuroimage 161, 80–93 (2017).

44. Gordon, E. M. et al. Precision Functional Mapping of Individual Human Brains. Neuron 95, 791–807.e7 (2017).

45. Cui, Z. et al. Individual Variation in Functional Topography of Association Networks in Youth. Neuron 106, 340–353.e8 (2020).

46. Laumann, T. O. et al. Functional System and Areal Organization of a Highly Sampled Individual Human Brain. Neuron 87, 657–670 (2015).

47. Kragel, P. A., Han, X., Kraynak, T., Gianaros, P. J. & Wager, T. D. fMRI can be highly reliable, but it depends on what you measure. doi:10.31234/osf.io/9eaxk.

48. Fox, M. D. Mapping Symptoms to Brain Networks with the Human Connectome. New England Journal of Medicine vol. 379 2237–2245 (2018).

49. Wilson, B. M., Harris, C. R. & Wixted, J. T. Science is not a signal detection problem. Proc. Natl. Acad. Sci. U. S. A. 117, 5559–5567 (2020).

50. Power, J. D., Barnes, K. A., Snyder, A. Z., Schlaggar, B. L. & Petersen, S. E. Spurious but systematic correlations in functional connectivity MRI networks arise from subject motion. Neuroimage 59, 2142–2154 (2012).

51. Power, J. D. et al. Methods to detect, characterize, and remove motion artifact in resting state fMRI. Neuroimage 84, 320–341 (2014).

52. Casey, B. J. et al. The Adolescent Brain Cognitive Development (ABCD) study: Imaging acquisition across 21 sites. Dev. Cogn. Neurosci. 32, 43–54 (2018).

53. Glasser, M. F. et al. The minimal preprocessing pipelines for the Human Connectome Project. Neuroimage 80, 105–124 (2013).

54. Avants, B. B. et al. A reproducible evaluation of ANTs similarity metric performance in brain image registration. Neuroimage 54, 2033–2044 (2011).

55. Jenkinson, M., Beckmann, C. F., Behrens, T. E. J., Woolrich, M. W. & Smith, S. M. FSL. Neuroimage 62, 782–790 (2012).

56. Smith, S. M. et al. Advances in functional and structural MR image analysis and implementation as FSL. Neuroimage 23 Suppl 1, S208–19 (2004).

57. Marcus, D. S. et al. Informatics and data mining tools and strategies for the human connectome project. Front. Neuroinform. 5, 4 (2011).

58. Hallquist, M. N., Hwang, K. & Luna, B. The nuisance of nuisance regression: spectral misspecification in a common approach to resting-state fMRI preprocessing reintroduces noise and obscures functional connectivity. Neuroimage 82, 208–225 (2013).

59. Gordon, E. M. et al. Generation and Evaluation of a Cortical Area Parcellation from Resting-State Correlations. Cereb. Cortex 26, 288–303 (2016).

60. Seitzman, B. A. et al. A set of functionally-defined brain regions with improved representation of the subcortex and cerebellum. Neuroimage 206, 116290 (2020).

61. Barch, D. M. et al. Demographic, physical and mental health assessments in the adolescent brain and cognitive development study: Rationale and description. Dev. Cogn. Neurosci. 32, 55–66 (2018).

62. Rothman, K. Modern Epidemiology. (Lippincott Williams & Wilkins, 2016).

63. Van Essen, D. C. et al. The Human Connectome Project: a data acquisition perspective. Neuroimage 62, 2222–2231 (2012).

64. Nystrom, N. A., Levine, M. J., Roskies, R. Z. & Ray Scott, J. Bridges. Proceedings of the 2015 XSEDE Conference on Scientific Advancements Enabled by Enhanced Cyberinfrastructure - XSEDE’15 (2015) doi:10.1145/2792745.2792775.

65. Towns, J. et al. XSEDE: Accelerating Scientific Discovery. Computing in Science & Engineering vol. 16 62–74 (2014).

